# Navigating a fine balance: point-mutant cheater viruses disrupt the viral replication cycle

**DOI:** 10.1101/2024.09.18.613812

**Authors:** Moran Meir, Arielle Kahn, Carmel Farage, Yael Maoz, Noam Harel, Adi Ben Zvi, Shir Segev, Maria Volkov, Ravit Yahud, Uri Gophna, Adi Stern

## Abstract

Cheater viruses, alternatively denoted as defective interfering viruses, cannot replicate on their own yet replicate faster than the wild type (WT) when the two viruses coinfect the same cell. Cheaters must possess dual genetic features: a defect, which leads to their inability to infect cells on their own, and a selective advantage over WT during co-infection. Previously, we have discovered two point-mutant cheaters of the MS2 bacteriophage. Here, we set out to discover the possible repertoire of cheater MS2 viruses by performing experimental evolution at a very high multiplicity of infection (MOI). Our results revealed a third point-mutant cheater that arose in eight biological replicas. Each of the three cheaters disrupts the fine balance necessary for phage replication, in different ways that create a defect + advantage. We found that over time, the point mutant cheaters accumulate additional “helper” mutations, which alter other stages of the viral replication cycle, complementing the disruptions created by the original cheater. Intriguingly, cheater and helper mutations almost always reside in very close proximity on the genome. This region encodes for multiple functions: overlapping reading frames as well as overlapping RNA structures critical for transitioning from one stage to another in the viral replication cycle. This region of overlap explains the dual functions of cheaters, as one mutation can have pleiotropic effects. Overall, these findings underscore how viruses, whose dense genomes often have overlapping functions, can easily evolve point-mutant cheaters, and how cheaters can evolve to alter the intricate balance of the viral replication cycle.

## Introduction

Viruses are ultimate parasites that rely on cellular infection to complete their replication cycle. In recent years, there is an increasing interest in what happens when two or more viruses infect the same cell. In some cases, the co-infecting viruses may be of very different genetic backgrounds, and this may lead to recombination or reassortment, processes that have a dramatic impact on the evolution of many viruses. Alternatively, co-infection may occur between similar variants of the same virus, leading to an ecological interaction between the two co-infecting viruses, which can be classified as cooperative or antagonistic (Segredo-Otero & Sanjuán, 2022), concepts explored within the emerging field of “sociovirology” (Altan-Bonnet & Chen, 2015; DaPalma et al., 2010; Díaz-Muñoz et al., 2017; Sanjuan & Thoulouze, 2019; Santiana et al., 2018).

Defective virus genomes that can thrive in the presence of co-infecting wild-type (WT) viruses, also known as defective interfering particles or cheater viruses, are attracting increasing interest. As we will go on later to define in the context of this paper, we will mainly use the term cheater viruses. Often, cheater virus genomes are truncated, lack essential genes, and are structurally distinct from the WT genomes. They rely on the co-infecting WT genomes to supply their gene products, also denoted as “public goods” (Chao & Elena, 2017; Ciota et al., 2012; Díaz-Muñoz et al., 2017); they furthermore subvert resources from the WT genomes, thus negatively impacting the replicative capacity of WT. Recent interest in cheater viruses has grown due to their potential therapeutic applications and a growing understanding on their potential effects on evolutionary processes.

On the therapeutic side, by competing for resources with pathogenic WT viruses, cheater viruses can reduce the infectious load and stimulate both adaptive and innate immune responses (Bdeir et al., 2019; Brennan & Sun, 2024; Dimmock & Easton, 2014; Genoyer & López, 2019; Leeks, Bono, et al., 2023; Rast et al., 2016; Tanner et al., 2019; Xu et al., 2017). There is a growing interest in the use of therapeutic interfering particles that are effectively cheater viruses (Chaturvedi et al., 2021; Pelz et al., 2021; Rezelj et al., 2021), and this has stimulated increased research in the field.

Viral cheating also has an impact on both short-term and long-term evolutionary dynamics. First, natural selection may be reduced in the presence of cheaters through the presence of defective cheaters or defective genomes that may thrive during co-infection. Cheaters may also lead to a collapse in the viral population, with implications on effective viral population size and weaker selection (Leeks et al., 2021; Singhal & Turner, 2021). Cheater viruses may impose selection on the WT viruses to avoid invasion by cheaters, e.g., via superinfection exclusion (Hunter & Fusco, 2022), and may even impose selection for new genomic forms, as has been suggested in the case of multipartite viruses (Leeks, Young, et al., 2023). Finally, cheaters add to the viral population an additional layer of genetic diversity that may lead to phenotypic novelty, similar to other selfish elements (e.g., transposons) that sometimes undergo exaptation and evolve to take on new functions (Fueyo et al., 2022; Hall et al., 2020).

We have previously discovered two point-mutant cheater viruses during experimental evolution of the MS2 bacteriophage, highlighting that cheating can occur not only via large genomic and structural variations, but via a mere point mutation (Meir et al., 2020). We hereby define a cheater virus as a virus bearing two traits: (i) a defect that reduces its ability to replicate on its own as compared to WT viruses, and (ii) a fitness advantage over WT during co-infection of cells with both WT and the cheater virus (Leeks et al., 2021). The second condition is similar to a fitness cost imposed on the WT (Ghoul et al., 2014); yet the definition herein allows to focus only on cheater characteristics.

MS2 is a model virus that has been studied in depth, due to its applications in biotechnology and the pioneering insights it has allowed into genome function and the viral replication cycle (Dang et al., 2023; Naskalska & Heddle, 2024). Its replication cycle consists of four main stages: (I) Entry, (II) Replication, (III) Packaging, (IV) Lysis/Exit (Fig. 1A) (Rolfsson et al., 2016; Van Duin & Tsareva, 2006). Each of these four stages is controlled by one of the four proteins the virus encodes for, respectively: (I) maturation (also known as A-protein), (II) replicase, (III) coat, (IV) lysis (Fig. 1B). Of note, the latter three viral protein products can be seen as “public goods” that can be utilized by cheaters during co-infection. Transitions between the four stages of the viral life cycle are tightly controlled by RNA structures in the virus genome that overlap the viral open reading frames (ORFs). The long-range Min-Jou (MJ) structure impacts replicase translation and hence genome replication (Van Himbergen et al., 1993). It has further been shown that the upstream coat terminator (CT) structure, residing immediately upstream the MJ, also impacts genome replication. Binding of coat protein to the TR loop structure initiates packaging of the virus (Johansson et al., 1997). Expression of lysis protein, and hence lysis of cells and exit of viruses, is controlled by the lysis hairpin (LH) loop (Fig. 1, Fig. S1) (Betancourt, 2009; Van Himbergen et al., 1993).

**Figure 1.**
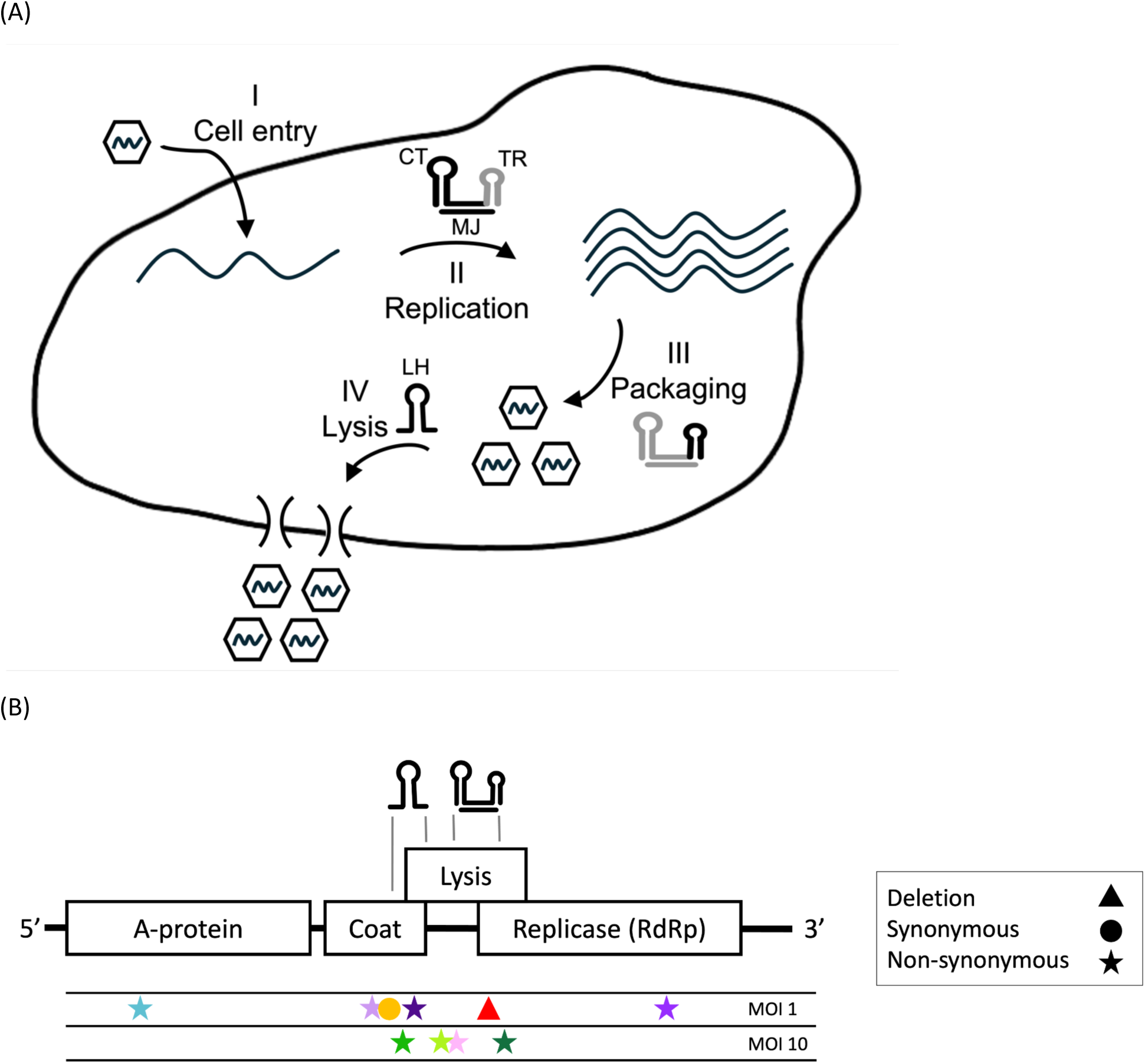
Scheme of MS2 replication cycle and genome. (A) A simplified schematic of the replication cycle of MS2 that is composed of four main stages: (I) Viral entry, which includes attachment to the host cell, and entry of viral RNA into the host cell, (II) Replication of positive sense RNA genomes (via negative strand intermediates – not shown), (III) Packaging and assembly of viral genomes into virions, (IV) Lysis of cells and exit of virions. All four stages are mediated through translation of a relevant viral protein and recruitment of host factors as well (not shown). Transitions between different stages are mediated through the illustrated viral RNA structures (main text). As the CT, MJ, and TR loop structures are adjacent, the relevant structure is highlighted in black versus grey (stages II and III). (B) A schematic of the MS2 genome with its four open reading frames, each of which mediates a different stage of the replication cycle (main text). RNA secondary structures are illustrated above the genome. The MJ structure is a long-range structure that spans beyond the boundaries of the illustration. Below the genome is a summary of all the major mutations found herein and previously in the context of high MOI serial passaging.

Briefly, the two MS2 point-mutant cheaters we previously found (Meir et al., 2020) consisted of a point deletion (Δ1764) and a synonymous mutation (A1664G) that impacts the LH structure. We were able to show that Δ1764 was unable to replicate on its own yet bore an advantage during co-infection via enhanced RNA genome packaging. Conversely, A1664G was defective on its own due to reduced lysis protein translation (producing tiny plaques) yet bore an advantage during co-infection likely via enhanced packaging as well. We denoted the phenotype of Δ1764 as full cheating since its fitness was zero on its own, and the phenotype of A1664G as semi-cheating, since its fitness was higher than zero on its own.

In this work, we set out to more systematically characterize the repertoire of cheater MS2 viruses and their evolution. A key term that impacts the ability of cheaters to emerge and thrive is that of multiplicity of infection (MOI), the ratio between the number of infectious virus as assessed by plaque assay and the number of bacterial cells. We focused first on a very high MOI of 10, where co-infections occur at a high rate during each passage. We further explored longer-term evolution of our previously discovered cheaters that were evolved at an MOI of 1. Notably, when cheaters emerge, they lead to an effective increase in the MOI (they are not counted during plaque assay that is used to determine MOI); thus, the effective MOI increases as cheater frequencies increase.

One of our goals was to test whether cheaters can correct their defect via compensatory mutations that corrected the defect. We found that different cheaters found in both experiments gained specific sets of additional “helper” mutations on their genomes, yet these did not correct the defect of the original cheater mutations. We tested the impact of these helper mutations and found that they tend to alter another one of the four stages of the replication cycle, one not necessarily overlapping that of the defect. We revealed that both cheater mutations and helper mutations tend to reside in a very specific region of the genome (Fig. 1B) with multiple overlapping functions. We propose a “fine balance model” whereby cheaters occur at regions of overlapping functions, disrupt one stage of the replication cycle and then evolve to alter a different stage, while overall maintaining high fitness during co-infection.

## Results

### Parallel evolution during serial passaging at MOI of 10

To test if additional point mutant cheaters emerge under conditions of higher MOI, we performed serial passaging at an MOI of 10. Passaging was performed across three different founder populations, resulting in eight biological replicas. The large number of replicas was performed to ensure that founding populations do not affect the biological outcome during passaging (Methods). Strikingly, we observed parallelism across all of the eight replicas and revealed that the same set of five mutations increased in frequency in parallel across all replicas (Fig. 2A), a strong signal of positive selection. The highest frequency was obtained for mutation A1744G, reaching a frequency of up to 75%. For simplicity, we will denote this mutation as A1744G-pink (Fig. 2). One of the five mutations observed was A1664G, the semi-cheater we have previously reported, which we henceforth refer to as A1664G-orange. Three additional mutations were also repeatedly observed: G1688U, C1735U, and U1829C, which we will denote as green helper mutations.

**Figure 2.**
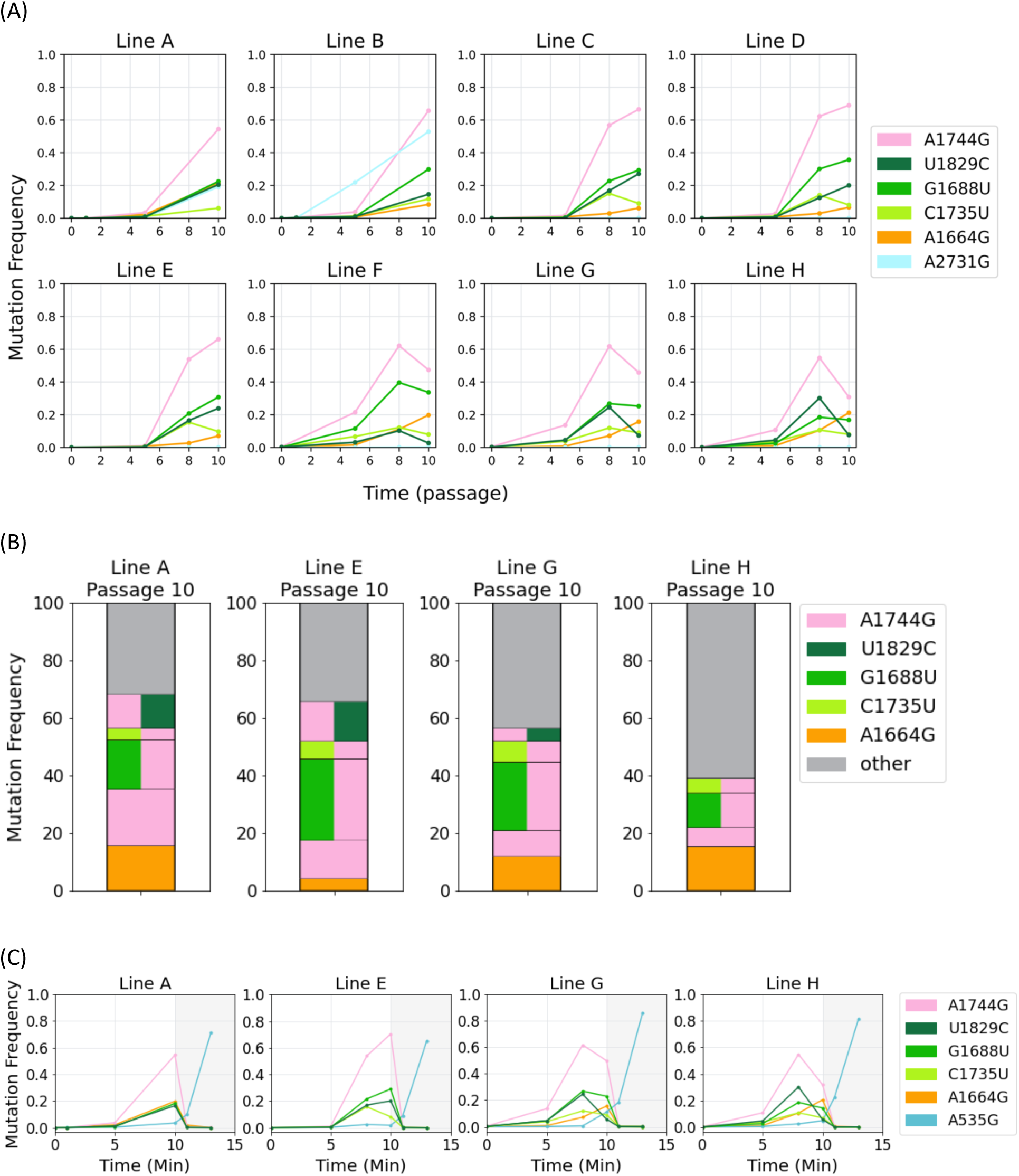
Parallel evolution during MOI=10 passaging. (A) Trajectories of mutations from eight replicas of serial passaging performed at an MOI of 10 reveal strong parallelism. Shown are mutations exceeding 10% in at least two or more replicas. (B) Results of full haplotypes from four p10 populations (lines A,E,G,H), derived from synthetic long read sequencing. Each horizontal block represents a haplotype (combination of mutations), and height is proportional to frequency of the haplotype. Only haplotypes exceeding 3% are displayed to allow a view of mutations observed in (A). (C) Similar to (A), with the addition of three terminal passages performed at a low MOI of 0.01. Passaging was reinitiated from a resuscitated passage 9 in three lines (A,E,G,H), and three additional passages (11-13; grey background) were performed at an MOI of 0.01. Shown are mutations exceeding 10%. See Fig. S5 for mutations at lower frequencies from (A) and (C).

We next used synthetic long read sequencing (Methods) to determine which of the five mutations we discovered reside on the same genome. The results showed a very consistent pattern: A1744G-pink was either on its own, or with exactly one additional green helper mutation (G1688U, C1735U, or U1829C). These latter green helper mutations were almost never found on their own, were never together with one another on the same genome and were never found with the semi-cheater A1664G-orange (Fig. 2B, Fig. S2).

To further explore the repertoire of cheating in MS2, we used the long-read sequencing data to search for genomes with large deletions, a hallmark of cheaters in other viruses (Pelz et al., 2021; Secor & Dandekar, 2020; Wu et al., 2022). We noted the presence of shorter genomic fragments, spanning either a few hundred bases or around 1800 nucleotides (Fig. S3). However, their frequency reached at most 0.5%, suggesting that they are not potent cheaters with a large advantage.

### A point-mutant cheater A1744G-pink is inferred

We set out to test whether the newly discovered mutations A1744G-pink and the green helper mutations are simply *bona fide* adaptive mutations or represent cheater viruses. To this end, we performed three additional passages using four p10 populations (lines A, E, G, H), at an MOI of 0.01. These conditions do not allow for co-infection and thus allow propagation of *bona fide* adaptive mutations but not of cheaters. Our results showed that A1744G-pink and the green helper mutations decreased to a frequency of zero. On the other hand, we observed an increase in the frequency of A535G-turquoise. This is a mutation we have previously shown to be a *bona-fide* adaptive mutation, supported by the results herein (Fig. 2). A535G-turquoise creates a non-synonymous mutation in the A-protein, thus likely allowing more rapid entry into cells.

We also sequenced a set of thirty-two plaques from a late passage and included plaques of various sizes (Methods). Our aim was to test if our newly discovered cheaters are full cheaters or semi-cheaters that can create smaller plaques. We did not observe any of the four mutations in any of the plaques (Fig. S4). Altogether, our results suggest that A1744G-pink is the driver mutation here that serves as a full point-cheater mutation. We propose that the green helper mutations, which are observed in all 8/8 replicas, provide some type of added benefit to the cheater, as explored below.

### Mechanism of pink cheating and corrections by green helpers

Per our definition, cheaters bear both a defect during singular infection, as well as an advantage during co-infection. We set out to infer these two traits for A1744G-pink. This was challenging since this mutation creates three potential phenotypes (Table 1): (a) a non-synonymous mutation at the lysis protein (K23E), previously shown to negatively impact lysis of cells (Chamakura et al., 2017) (b) alteration of the MJ RNA structure that controls translation of the replicase protein, thus possibly reducing or enhancing replication, and (c) alteration of the TR loop RNA structure, which halts replicase translation and initiates packaging, thus possibly reducing or enhancing packaging.

**Table 1.** Inferred phenotypic effects of A1744G-pink and green helper mutations G1668U, C1735U, U1829C.

| Mutation | Type | Affected protein/structure | Inferred phenotype | Replication cycle stage (Fig. 1)<sup>a</sup> |
| --- | --- | --- | --- | --- |
| A1744G | Non-synonymous | Lysis (K23E) | Reduced lysis expression (Chamakura et al., 2017) | IV |
|  | RNA structure | MJ | Alteration of the MJ RNA structure → reduction or enhancement of replicase translation (Fig. S1) | II |
|  | RNA structure | TR | Alteration of the TR loop → reduction or enhancement of packaging (Fig. S1) | III |
| G1668U | Synonymous | Coat | -- | -- |
|  | RNA structure | LH | Increased stability of RNA hairpin structure → decreased ribosome binding → reduced lysis expression (Berkhout et al., 1987; Schmidt et al., 1987) | IV |
|  | Non-synonymous | Lysis (R4L) | Unknown | IV |
| C1735U | Non-synonymous | Lysis (R20W) | Reduced lysis expression (Chamakura et al., 2017) | IV |
| U1829C | Synonymous | Replicase | -- | -- |
|  | Non-synonymous | Lysis (F51S) | Reduced lysis activity (Chamakura et al., 2017) | IV |

<sup>a</sup> I – cell entry, II – replication, III – packaging, IV - lysis

We thus set out to test the effects of A1744G-pink on replication using an assay of intracellular replication. We sequenced genomes from pellet (intracellular genomes) at 15, 30,45,60, 90 minutes, as well as overnight, using populations of p8-D. This was compared to supernatant (virion) sequencing at time points zero and overnight (ON). Two such assays were performed, one at an MOI of 0.01 (Fig. 3A) and one at an MOI of 10 (Fig. 3B). Importantly, the low MOI assay allowed us to infer how each of the possible genotypes behaves during singular infection, since such low MOI ensure each cell is infected by exactly one genotype. Conversely, the high MOI intracellular assay allowed us to infer the behaviour of genotypes under the conditions of the original serial passaging experiment.

**Figure 3.**
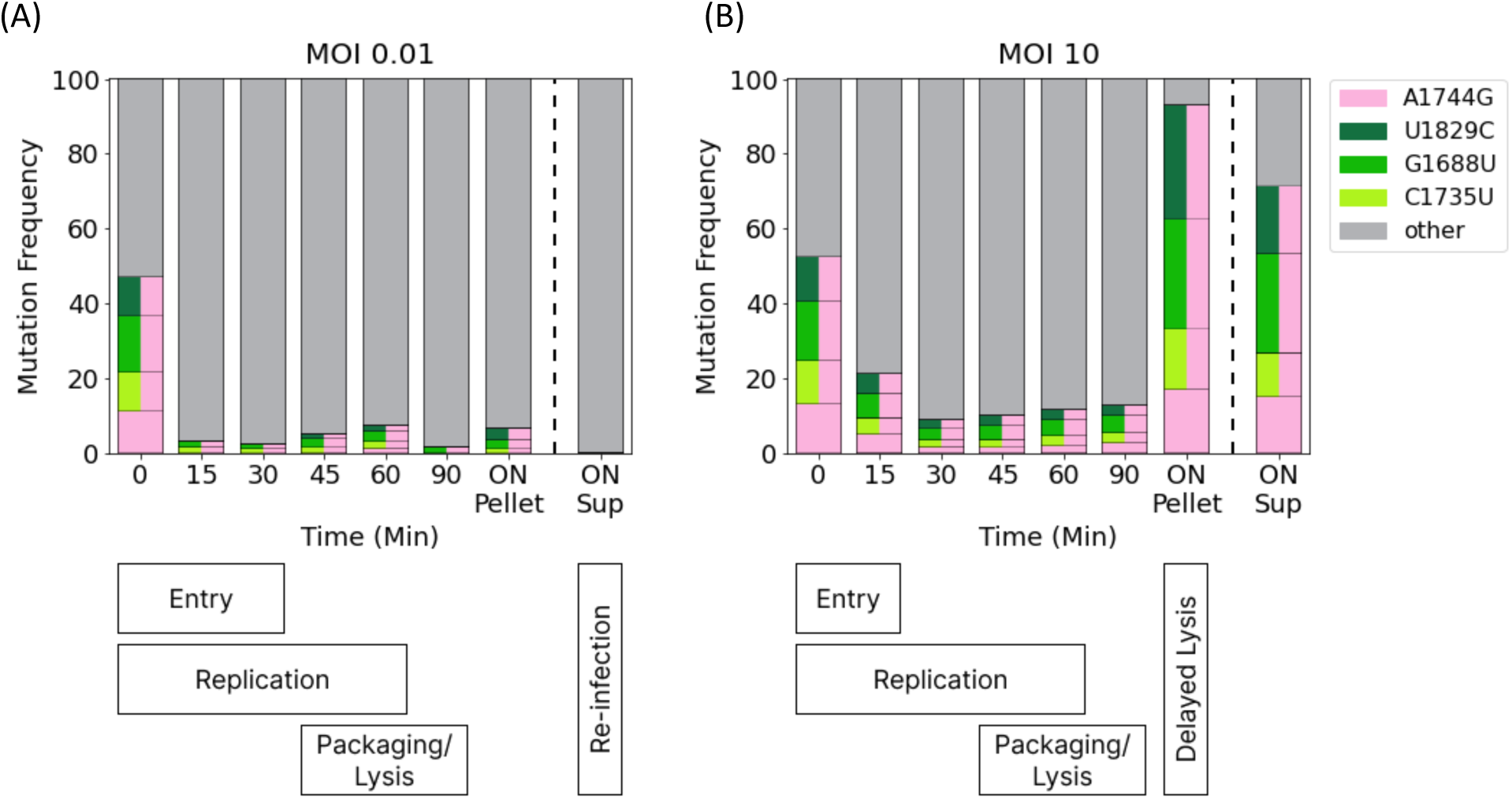
Intracellular assay of replication for A1744G-pink and green helper mutations. The frequencies of the various genotypes are shown across various time points of the viral replication cycle from genomes isolated from inside infected cells (Pellet). Time points zero and ON-Sup represent sequencing of virions from supernatant at the start and end of a passage. Only genotype frequencies initially higher than 3% are displayed. Genotype frequencies were inferred from Fig. 2B (Methods). (A) Results from the intracellular assay at MOI of 0.01, (B), Results from the intracellular assay at MOI of 10. The lower panels illustrates timing of entry, replication, and packaging/lysis based on plaque assay and qRT-PCR measurements (Fig. S6). Of note, replication is evident as of 7 mins post infection (Fig. S6), but for simplicity is illustrated as starting at time zero. Presence of mutants in overnight (ON) Pellet at high MOI suggests delayed lysis, whereas increased presence of mutants in ON Sup suggests increased packaging. Differences between low and high MOI timing stem from delayed entry at low MOI, as well as an additional replication cycle (reinfection) at low MOI.

The frequency of the A1744G-pink and green helper mutations all dramatically declined between 0 and 15 minutes at both MOIs, suggesting either a replication disadvantage (stage II) or a cell entry disadvantage (stage I) associated with these mutations. Since A1744G-pink was not inferred to impact entry in any way (Table 1), and neither were the green helper mutations, we conclude that A1744G likely leads to a defect in replication due to its impact on the MJ structure. This defect is probably what led to no A1744G-pink in overnight extracellular/sup populations at MOI of 0.01. The defect is compensated at higher MOI by co-infection and use of WT replicase, and hence less of a decline of A1774G-pink genomes is observed at high MOI at 15 minutes.

We noted a strong increase in frequencies of A1744G-pink (and green helper mutations) in the overnight pellet populations tested at MOI=10 (Fig. 3B), with the largest increase in A1744G-pink combined with either U1829C-green or G1668U-green. This result suggests either a replication advantage (which we ruled out above), or delayed lysis, allowing more genomes to remain intracellular. We see only minor frequencies of A1744G-pink in overnight pellet populations at an MOI of 0.01, since there is likely another round/s of replication that releases WT viruses overnight. Given that there is an inferred defect in lysis created by all of the green mutations (as well as for A1744G-pink) (Table 1) (Chamakura et al., 2017), we suggest that they all contribute to delayed lysis. Under conditions of high MOI (no more cells to re-infect) and under conditions of defective replication by A1744G-pink, delayed lysis may be an advantage as it provides additional time for genome replication of the cheaters.

What if so, is the advantage of A1744G-pink during high MOI that leads to the increase in A1744G-pink during serial passaging, also reflected in the overnight sup populations at high MOI? The results suggest an advantage that must be post-replication. Given the potential effects of A1744G-pink on the TR loop structure (Table 1), we suggest that this advantage is in packaging. However, an alternative non-mutually exclusive explanation is that as described above, delayed lysis in itself provides an advantage under conditions where there is not an additional replication cycle: during the time the viruses are “trapped”, or partially trapped, in the cells, replication continues. If lysis is not completely defective, some of these viruses may exit the cells during the long wait overnight. To summarize, we conclude that A1744G-pink is a replication-defective cheater that uses WT replicases as “public goods” during co-infection. A1744G-pink likely has an advantage in packaging, and a possible defect in lysis as well. Green helper mutations that emerge on the A1774G-pink background serve to delay lysis and allow more time for replication.

### Revisiting serial passaging at MOI of 1 reveals more helper mutations

In parallel to our serial passaging experiments at an MOI of 10, we had also previously performed serial passaging at MOI of 1 in two replicas, and originally, we observed competition between the two cheaters (A1664G-orange and Δ1764-red), with A1664G-orange emerging as a “winner” (Meir et al., 2020). We continued the passaging at an MOI of 1 till passage 30 (Fig. 4), originally with the aim of testing (a) whether a steady state would be obtained, and (b) whether the semi-cheater A1664G-orange would “correct” its defect.

**Figure 4.**
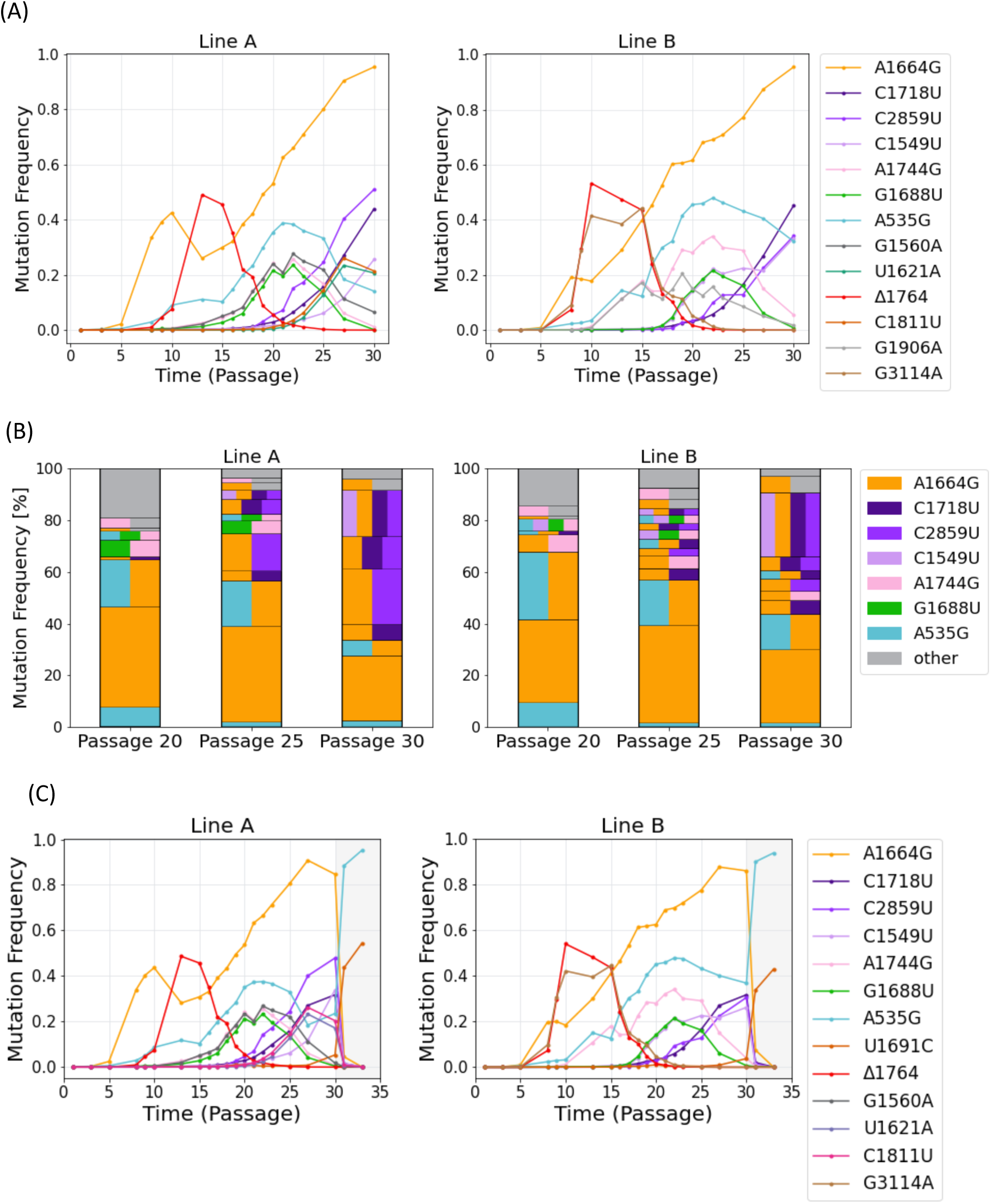
Parallel evolution during advanced MOI=1 passaging. (A) Trajectories of mutations from two replicas of serial passaging performed at an MOI of 1, initiated in (Meir et al., 2020) and continued herein from a resuscitated p23 population. Due to the large number of mutations that emerge, shown are mutations with *f*≥20% in any replica. (B) Results of full haplotypes from different passages from both lines. Haplotypes structures as described in Fig. 2B. Only haplotypes with *f*≥3% at one or more time points are displayed (A). (C) Similar to (A), with the addition of three terminal passages performed at low MOI of 0.01. Passaging was reinitiated from a resuscitated p30 in both lines, and three additional passages (11-13; grey background) were performed at an MOI of 0.01. Shown are mutations with *f*≥20%. See Fig. S8 for mutations at lower frequencies from (A) and (C).

Our results showed that A1664G-orange continued to increase in frequency, attaining a frequency of over 90%. Once again performing long-read synthetic sequencing allowed us to resolve haplotypes. Interestingly, in both replicas we noted a temporary rise and demise of the A1744G-pink, occasionally with one of the green helper mutations (G1688U) and occasionally with A1664G-orange. We also observed the rise and decline of A1664G-orange combined with A535G-turquoise (Fig. 4B). Finally, we observed a set of three so-called purple helper mutations that began to increase in frequency as of passage 20, which all resided on genomes together with A1664G-orange. As opposed to the green helpers described above, we did observe combinations of the purple mutations with each other. The decline of all combinations of mutations, versus the rise of combinations of A1664G-orange and the purple mutations, led us to conclude that the latter combinations are the most fit in the conditions of the passaging.

We noted that two of the three purple mutations (C1718U and C1549U) reside in the same region where there are multiple overlapping functions (Fig. 1B), rendering it challenging to infer their function (Table 2). The third, C2859U, creates a non-synonymous mutation in the replicase gene, which resides one residue upstream a functional motif of the encoded protein (Fig. S7), lending a first clue as to the combined function of these mutations. In fact, in principle, all three mutations are inferred to impact replication (Table 2), as tested below.

**Table 2.** Inferred phenotypic effects of A1664G-orange and purple helper mutations C1549U, C1718U, C2859T.

| Mutation | Type | Affected protein/ structure | Inferred phenotype | Replication cycle stage (Fig. 1)<sup>a</sup> |
| --- | --- | --- | --- | --- |
| A1664G | Synonymous | Coat | -- | -- |
|  | RNA structure | LH | Reduced lysis translation (Meir et al., 2020). | IV |
| C1549U | Non-synonymous | Coat (T72I) | Reduced coat protein binding to packaged RNA genome, may alter virion structure and/or alter RNA release during cell entry. | I |
|  |  |  | Reduced coat protein binding to TR loop, may delay packaging and increase replication (Lago et al., 1998; Stonehouse et al., 1996). | II/III |
| C1718U | Synonymous | Coat | -- | -- |
|  | RNA structure | Coat terminator | Destabilizes coat terminator (Fig. S1) → may increase replicase translation and hence replication (Van Himbergen et al., 1993). | II |
|  | Non-synonymous | Lysis (A14V) | Unknown | IV |
| C2859T | Non-synonymous | Replicase (R367C) | Adjacent to predicted replicase motif E (RNA binding) (Fig. S7). May promote or decrease replication. | II |

<sup>a</sup> I – cell entry, II – replication, III – packaging, IV – lysis

### Does the high frequency of A1664G-orange imply a loss of cheating phenotype?

We considered it possible that the A1664G-orange semi-cheater “corrected” its defect via the helper mutations, while retaining its advantage. To test this, we resuscitated the p30 samples and performed additional passaging at a low MOI of 0.01 that does not allow for co-infection (Fig. 4C). This assay revealed that the frequency of A1664G-orange and that of all the helper mutations – crashed to zero. Two other mutations increased in frequency, A535-turquoise, described above, and U1691C, a mutation we have previously observed as adaptive under low MOI conditions probably via more rapid lysis of cells (Caspi et al., 2023). We conclude that the A1664G-orange semi-cheaters remain as cheaters during the entire duration of passaging in the MOI=1 experiment.

### Mechanism of purple helper mutations

We set out to test whether we can infer the function of the purple helper mutations. To this end, we once again performed an assay of intracellular replication, and sequenced genomes at 0, 15, 30,45,60, 90 minutes, and overnight, using populations of p28-B (Fig. 5). Importantly, as described above, MOI of 0.01 allows us to assume each cell is infected by one genotype, allowing us to make inferences about effects of each genotype on its own. MOI of 1 allows us to obtain a picture of what occurs during co-infection at the same conditions as the original experiment was performed.

**Figure 5.**
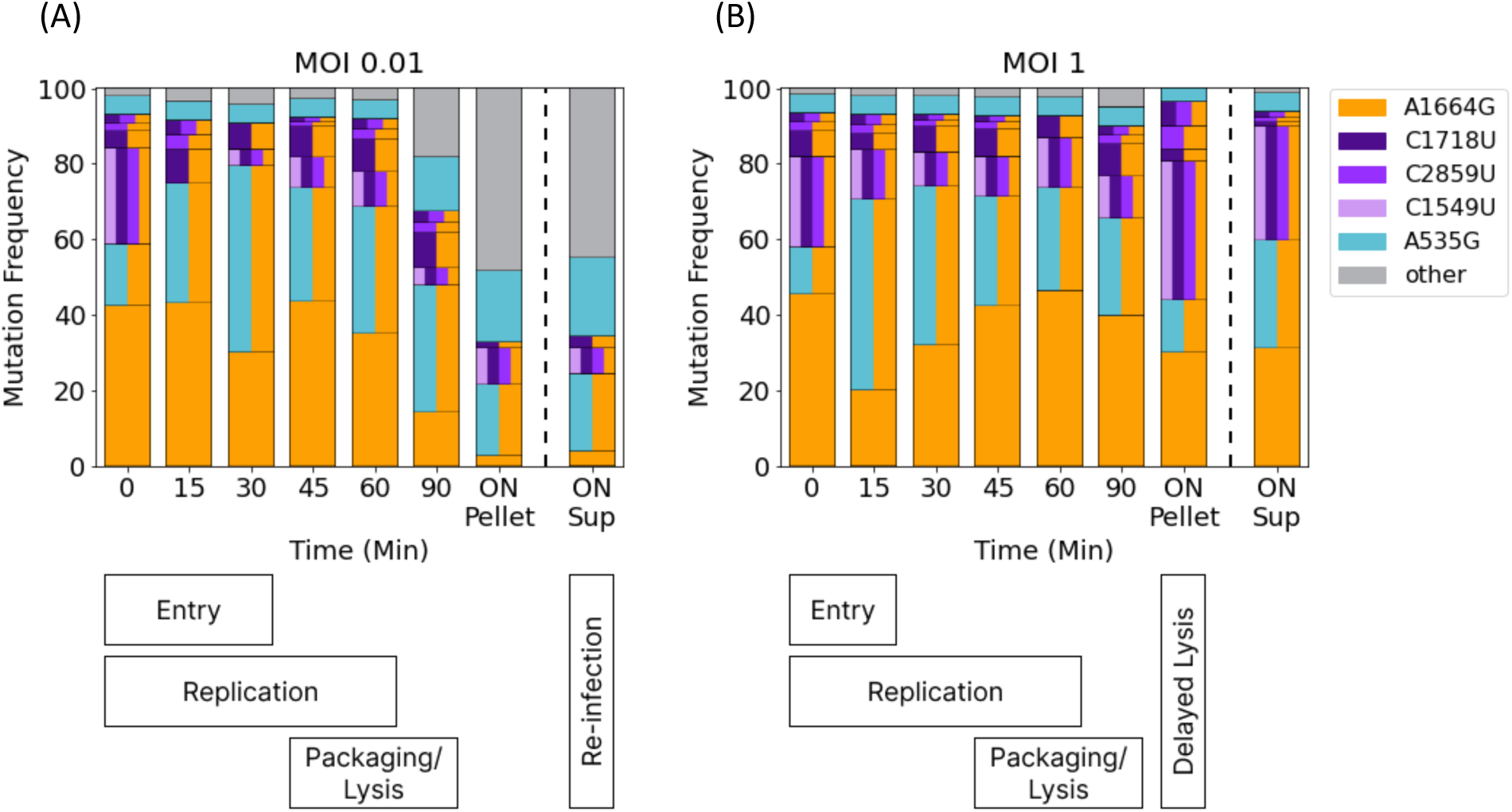
Intracellular assay of replication for A1664G-orange and additional mutations. The frequencies of the various genotypes are shown across various time point of the viral replication cycle from genomes isolated from inside infected cells (Pellet). Time points zero and ON-Sup represent sequencing of virions from supernatant at the start and end of a passage. Only genotype frequencies initially higher than 5% are displayed. Genotype frequencies were inferred from Fig. 4B (Methods). (A) Results from intracellular assay at MOI of 0.01, (B), Results from intracellular assay at MOI of 10. The lower panel illustrates timing of entry, replication, and packaging/lysis (see Fig. 3 legend).

First, as we have previously shown (Meir et al., 2020), A1664G-orange was not impacted in replication capacity, as is evident from the steady proportion of A1664G-orange during the entire first sixty minutes. During overnight at MOI of 0.01, we observed a decrease in A1664G-orange in extracellular virions down to a frequency of ∼30%, reiterating its ability to replicate on its own that is inferior to WT (semi-cheater phenotype).

When observing A1664G-orange in the MOI=1 intracellular assay, we saw a more or less steady proportion of A1664G-orange across all time points, with a slight increase in intracellular overnight genomes. The latter reiterated the defect in lysis we have previously shown (Meir et al., 2020). In general, the population used here for the intracellular replication assay was much more complex (many more mutations and many more combinations) than the population used for the intracellular assay described earlier for A1744G-pink. This necessitated us to develop a simple model of intracellular replication at low MOI, with fitness parameters *w* for three of the four replication cycle stages where we infer an impact by the various mutations: entry, replication, and lysis. Lysis effectively captures the two last stages together of packaging and exit. An additional nuisance parameter was added to allow for re-infection during low MOI (Methods).

Each parameter was assumed to affect the frequency of genotypes at a specific stage of intracellular replication. Thus, for example, a mutant with more rapid replication than WT, would be associated with a parameter *w_r_* > 1 that allows for an increase in the frequency of this mutant at times points where replication occurs (15 through 60 mins; Methods, Fig. S6). The model allowed us to simulate the trajectory of intracellular frequencies of a mutant given its associated fitness values, and to fit this trajectory to the observations depicted in Fig. 5. We next used an approximate Bayesian computation (ABC) approach to infer parameters for the genotypes shown in Fig. 5.

To avoid over-parameterization, we combined all purple mutations together and referred to them as the “purple combo”. Our inferences suggested that the purple combo negatively impacted cell entry (posterior probability > 0.99), but also had an advantage during intracellular replication (posterior probability > 0.98). We did not find evidence to support or rule out that the purple combo led to more delayed lysis (Fig. S9). These former results can be derived by observing the MOI=1 intracellular assay results: we saw a decrease in the purple combo from 0 to 15 minutes, supporting a cell entry defect, and an increase in purple combo over time that was particular evident in the overnight pellet sample, supporting an advantage in replication.

To summarize, we conclude that A1664G-orange is a lysis-defective cheater that uses WT lysis as “public goods” during co-infection. A1664G-orange likely has an advantage in packaging as ribosomes not translating lysis will create more coat protein. Purple helper mutations that emerge on the A1664G-orange background serve to increase the replication rate thus allowing for more lysis to be translated. The latter advantage become critical as the frequency of A1664G-orange increases and the probability of co-infection with WT decreases.

## Discussion

In this work, we set out to discover the repertoire of cheaters in the MS2 bacteriophage, and their subsequent evolution during high MOI conditions. Interestingly, as opposed to work in previous viruses, we did not note any structural variants and in particular did not note high frequencies of large deletions. It is possible that large deletion cheaters exist in MS2, but that they are outcompeted by the potent cheating of the point-mutant cheater viruses. Moreover, as packaging signals exist throughout the MS2 genome (Dai et al., 2017; Rolfsson et al., 2016), this may render large deletion cheating less effective.

It was intriguing to note that all the three cheaters we have found here and previously (Δ1764-red, A1664G-orange, A1744G-pink), and most of the so-called helper mutations that subsequently evolved on the background of two of these cheaters, resided in a region spanning less than 300 nucleotides (Fig. 1B), i.e., less than 10% of the genome. This region is the densest region in the MS2 genome in terms of genomic information: there are two ORFs, and a series of RNA structures that overlap one or two of these ORFs. Analyzing the evolutionary rate of this region amongst homologs of MS2, we find that this region is one of the more conserved regions, displaying for most part a low rate (Fig. S10). This is not surprising given the necessity to maintain the function of the two proteins and critical RNA structures that reside there. Yet, we find that cheaters specifically tend to evolve in this region. We suggest that this stems from the nature of point-mutant cheaters – they must both disrupt one function and gain an advantage of another function, and this is possible exactly at regions of overlapping functions. Cheaters “escape” the rigidity of selection operating at these regions due to their reliance on co-infection.

We go on to discuss the role of the helper mutations. In the case of the A1744G-pink mutation, all of the three additional green helper mutations were ones that were inferred to be deleterious to lysis production or function. As we discussed above, we propose that delaying lysis is actually advantageous when: (a) there are no additional hosts to infect at the end of each infection cycle (high MOI), and (b) when replication is impaired on the background of A1744G-pink, and then delayed lysis gives more time for replication to occur. Thus, the fine balance of the replication cycle is disrupted by A1744G-pink that leads to reduced replicase production (Table 1). This defect this is partially corrected by the green helper mutations that delay lysis and allow time for more replication (Fig. 6A).

**Figure 6.**
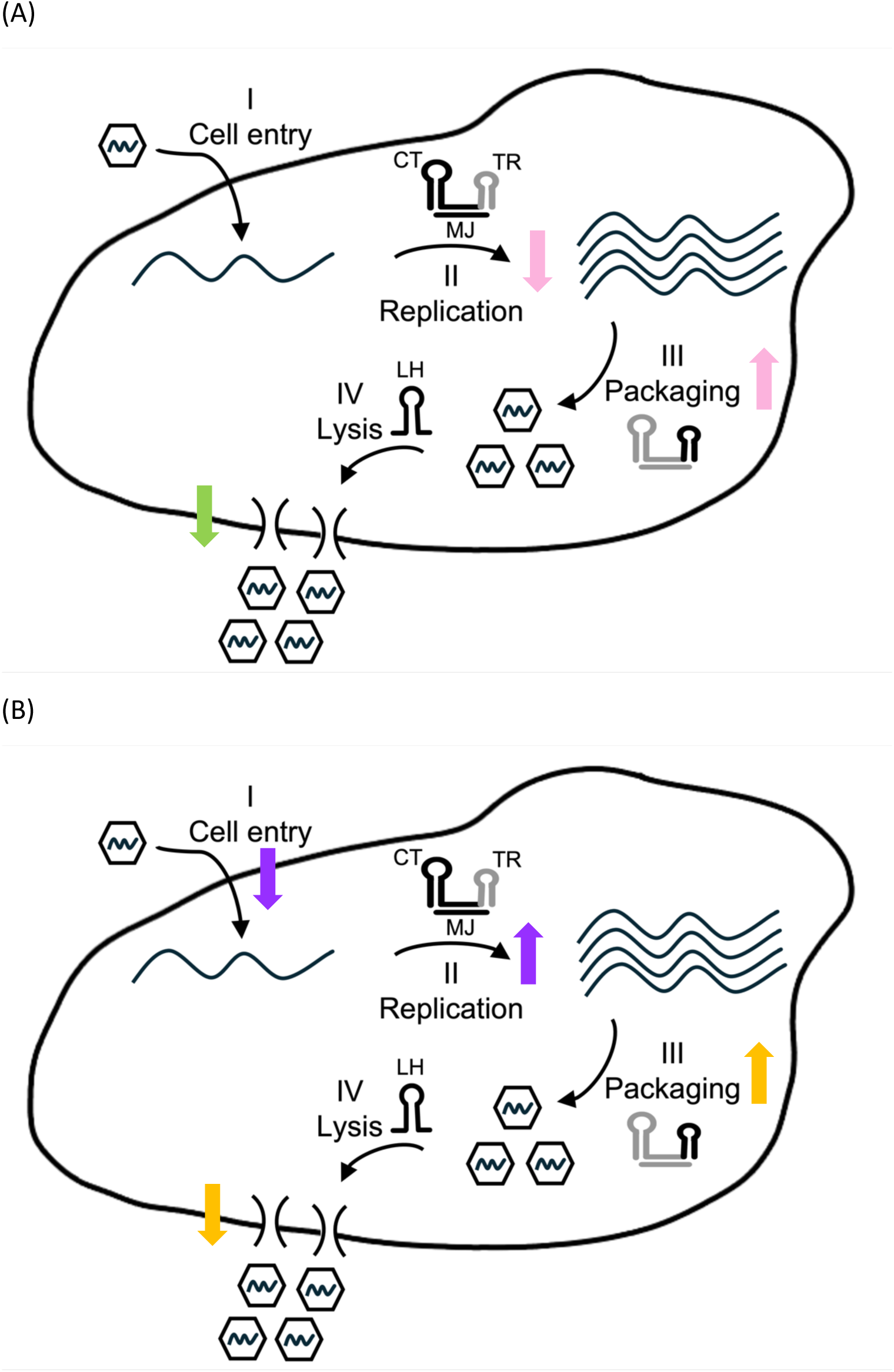
Fine balance model of viral replication cycle during viral cheating. Shown is the disruption of the cycle by cheaters and correction by helper mutations for the scenario of the A1744G-pink cheater (A) and the A1664G-orange semi-cheater (B). The four main stage of viral replication are shown as in Fig. 1A. Upwards or downwards colored arrows represent an increase or decrease in rates of a relevant stage based on the inferences deduced herein. The illustration describes a singular infection by a given genotype, and the number of phages that exits the cell is not modelled in the illustration.

If so, why does delaying lysis involve the particular three green mutations we observed, and why do we not see the green mutations together on the same genome? Overall, the MS2 lysis protein remains an enigma, and its mechanism of function has not been resolved. A recent study found it operates as an oligomeric complex (Mezhyrova et al., 2023). We suggest that lysis delay is achieved by mutants that create only a partial “poisoning” of the lysis oligomeric complexes that are composed of WT and mutants during co-infection. Accordingly, we suggest that double mutants, or other mutants of lysis (that are not one of the three we found), may cause a dominant-lethal phenotype – either through rapid lysis or no lysis at all.

We go on to discuss the advantage of the purple helper mutations on the background of A1664G-orange. Here, the situation is more complicated. One possibility is that all three purple mutations enhance replication (Fig. 6B). Under conditions of delayed lysis that A1664G-orange creates, this would be an advantage – more genome replication would lead to more lysis protein production that would compensate and allow for cell lysis. Notably, this is to some extent the mirror image scenario of what happens with A1744G-pink and its green helper mutations.

For C2859U-purple it is likely that replication is impacted, since the only phenotype this mutation is expected to create is a non-synonymous mutation in the replicase protein. It is also possible that replication is delayed by C1549U-purple and C1718U-purple. However, the latter two mutations may create other effects as well or on their own. C1718U-purple may additionally affect lysis or replication in an unknown way (Table 2). C1549U-purple was inferred to delay cell entry, and this seems like a clear-cut disadvantage. However, it may also allow for enhanced replication and this latter advantage may offset its disadvantage. Overall, an interesting observation we can deduct from the above, is that when a cheater disrupts one or more stages of the replication cycle, the subsequent mutations that emerge on the cheater background alter other stages of the replication cycle. Thus, there seems to be a fine balance of the viral replication cycle that can be shifted by “pulling the strings” of the various stages.

We would like to emphasize that although MS2 has a straightforward replication cycle, it serves as a model virus for many RNA viruses that undergo the same fundamental stages, albeit with additional layers of complexity. Moreover, it is possible that point-mutant cheaters that we find here to occur in regions with overlapping functions, may actually be prevalent and overlooked in many other RNA viruses that also tend to have regions with overlapping functions. It is intriguing to consider that cheaters may also serve as stepping stones in evolution, which allow viruses to overcome the stringent selection operating in such overlapping function regions. In other words, cheaters may evolve into cooperators, a concept that has been widely discussed under the umbrella of multilevel selection (Fiegna et al., 2006; Simon et al., 2013; Traulsen & Nowak, 2006).

This leads to the discussion on how prevalent high MOI conditions are in a natural setup. First, it has been shown that several viruses transit via collective infectious units (Leeks et al., 2019; Sanjuán, 2017), which is somewhat akin to higher MOI as it allows multiple virus genotypes to enter cells. More generally, MOI tends to cycle between high and low in most conditions, similar to predator prey cycles, and thus cheaters probably emerge and disappear cyclically. However, we believe it is possible that high MOI may be possible for a few replication cycles, as is observed during within-host evolution of many animal viruses (Felt et al., 2021; Vignuzzi & López, 2019). Over billions of years of evolution, cheaters will thus constantly emerge and evolve, potentially facilitating exploration of the fitness landscape in ways that could result in significant genetic innovations.

## Methods

### Stock preparation of MS2 phage

MS2 phage (from ATCC 15597-B1) and the F pilus expressing donor *E. coli* c-3000 were cultured in LB medium at 37°C with shaking overnight. Bacteria cells were removed by centrifugation at 4,000 rpm for 20 min at 25°C. The supernatant was filtered with a 0.22 μm filter (Stericup® Filter, EMD Millipore) to remove any remaining residues. The phage was then stored at 4°C. Typically, a stock concentration of 10^11^-10^12^ plaque forming units (PFU)/ml was obtained.

### MS2 plaques isolated

All the experiments started using a single WT MS2 plaque collected during plaque assay. A single MS2 plaque was collected with a sterile Pasteur pipette and transferred to an Eppendorf tube containing 1 ml of 0.85% NaCl. The tube was stored at 4°C overnight to facilitate viral particle release from the agarose. Next, 10 µl of chloroform was added, followed by vortexing and centrifugation for 3 minutes at 13,000 rpm at room temperature. The supernatant was then filtered through a 0.22 μm Minisart syringe filter (Sartorius) to remove residues. RNA was extracted from each plaque for whole genome deep sequencing, as detailed below. In addition, plaques in three different sizes (small, medium and large) were isolated from p8-D stock using the same method.

### Serial passaging

Clonal MS2 stocks were propagated from single plaques to ensure that the experiments will begin with a phage population as homogeneous as possible. Each clone served as the precursor for all evolutionary lines in each experiment. Serial passage experiments were conducted with two or three biological replicates, as followed. Three similar experiments were performed under the same conditions of MOI 10, Lines A to H. The first experiment included lines A and B, the second experiment included lines C to E and the third experiment included lines F to H. An additional experiment previously described (Meir et al., 2020) of serial passaging at MOI 1 was performed in two biological replicates, then denoted as lines 37A and 37B. The procedure for serial passaging was as follows: 100 ml cultures of naive *E. coli* c-3000 were grown at 37°C to an optical density of OD_600_ = 0.5, corresponding to approximately 10^8^ bacterial cells/ml (total of 10^10^ bacterial cells). For a multiplicity of infection (MOI) of 10, each passage was infected with 10 ml of 10^10^ phages (pfu) from the previous passage (total of 10^11^ pfu); for an MOI of 1, each passage was infected with 10 ml of 10^9^ phages from the previous passage (total of 10^10^ pfu), maintaining an MOI of about 10 and 1 pfu/cell respectively at the start of each passage. The cultures were incubated at 37°C for overnight with shaking, and the *E. coli* cells were subsequently removed by centrifugation. The supernatant was filtered through a 0.22 μm filter (Stericup® Filter, EMD Millipore) to eliminate any remaining residues. The resulting phage stocks were then stored at 4°C. Aliquots of these stocks were used to measure phage concentration via plaque assays, infect subsequent serial passages, isolate RNA for whole genome deep sequencing (as detailed below), and maintain frozen stocks of the evolving lines in 15% glycerol at −80°C.

For the experiments testing the effects of lowering the MOI, additional low MOI passages were conducted using the same protocol. This time, a frozen stock of either p30 line 37A MOI 1 and p30 line 37B MOI 1 or p10-C and p10-D was used as the starting point to test the effect of a lower MOI of 0.01 on the cheater population.

### Plaque assay

A plaque assay procedure was used to determine the concentration of the MS2 phages at each passage. The plaque assay was conducted according to the method described in (Meir et al., 2020).

### RNA Isolation

RNA was isolated using two methods based on the experiment:

1. Typically, RNA from all serial passages stocks as well as RNA from the supernatant of the replication assay was isolated using the QIAamp® viral RNA mini kit (Qiagen) according to the manufacturer’s instructions.
2. For the purpose of replication assays that test intracellular RNA, total RNA from *E. coli* cells (pellet) was isolated using the RNeasy Mini Kit (Qiagen) according to the manufacturer’s instructions.

### Illumina NGS library preparation

The MS2 RNA was reverse transcribed using SuperScript® 4 Reverse Transcriptase (Thermo Scientific), using the 3-R3 primer (Table S1). cDNA from the reverse transcription reaction was directly used as a template for the PCR amplification of the full MS2 genome in either three or seven overlapping fragments, and PCR reactions were performed using the Phusion High-Fidelity DNA-polymerase (Thermo Scientific) according to manufacturer instructions using the primers that are listed in Table S1. As quality control, we sequenced a set of given samples using either three or seven amplicons and the sequencing results were reproducible (not shown). The PCR products were purified using Wizard Gel and PCR Cleanup System (Promega). Purified amplicons were diluted and pooled in equimolar concentrations. To produce DNA libraries, the Illumina Nextera XT library preparation protocol and kit (FC-131-1096) were used according to manufacturer instructions. Libraries were sequenced on an Illumina Miseq or Nextseq using various reagent kits for paired end reads (MS-103-1002 Illumina, MS-102-2003 Illumina). All libraries were prepared in house and sequenced in the Genomics Research Unit at Tel-Aviv University (Israel), with the exception of the intracellular assay libraries that were prepared and sequenced by the Technion Genomics Center, Technion-Israel Institute of Technology, Haifa, (Israel).

### Read alignment and analysis of Illumina NGS

Illumina Miseq fastq files were initially processed using adapter removal and trimming. Adapter removal was performed using the Trimmomatic tool (Bolger et al., 2014) using ILLUMINACLIP for paired end sequencing with parameters: 2 seed mismatches, 30 palindrome clip threshold and 10 simple clip threshold. Trimming was performed with seqtk tool (https://github.com/lh3/seqtk) using trimfq to cut 30 bases from the beginning and end of each read. Analysis of the sequencing data was performed using the AccuNGS pipeline (Gelbart et al., 2020) with the parameters: minimal %ID=85, e-value threshold= 1E-07 and q score cutoff of 30. The pipeline uses BLAST (Altschul et al., 1990; McGinnis & Madden, 2004) to map the reads to the reference genome, searches for variants that appear on both overlapping reads and calls variants with a given Q-score threshold. Finally, it infers the frequency of each variant in the population. The reference genome is as described in (Meir et al., 2020). Libraries attained a minimum coverage of 100 reads per base. Positions at both ends of the genome (at the primer positions plus 5 adjacent positions) were removed from analysis because they had low coverage and higher variability.

### Library construction, sequencing and processing of synthetic long reads

RNA samples from passages p20, p25, and p30 from both lines (37A and 37B) at MOI 1 were sent to Element Biosciences (San Diego, CA, USA) for sequencing. The sequencing was performed using the LoopSeq RNA preparation kit following the manufacturer’s instructions (information available at elementbiosciences.com). The LoopSeq protocol (Callahan et al., 2021) utilizes unique molecular barcoding technology, which distributes barcodes evenly across a genome before fragmenting it into shorter pieces. These labeled fragments are then sequenced using short-read sequencing methods on the AVITI platforms, followed by the reconstruction of full-length genomes. The short-read raw data was processed through the Loop Genomics analytical pipeline, which handles low-quality base trimming, unique sample barcode demultiplexing, and synthetic long-read reconstruction. This process allows for de novo assembly of full-length genomes by rearranging short reads tagged with the same unique barcode.

We used BLAST (Altschul et al., 1990; McGinnis & Madden, 2004) to align the long reads obtained from LoopSeq with the MS2 reference sequence (Meir et al., 2020). The alignment was performed using the following parameters: --evalue 1e-07 --perc_identity 0.85 --task blastn --num_alignments 1000000 --dust no --soft_masking F. To ensure that only reads spanning the full alignment were considered, we filtered out any alignments that mapped more than once to the reference and those shorter than 3,500 nucleotides (98% of the reference genome length). Additionally, we excluded alignments that aligned to the minus strand, i.e., those that were the reverse complement of the genome.

### Intracellular replication assay

*E. coli* c-3000 cells were grown to an optical density of OD_600_=0.5 at 37°C. 100 ml of bacteria were infected with 10 ml of different viral populations: p28 line 37B at MOI 1 and p8-E, each one diluted to produce an MOI of 0.01, 1 or 10 pfu/cfu. The cultures were grown at 37°C with shaking. 1 ml volumes of the cultures were removed at 0, 15, 30, 45, 60, 90 minutes post infection, and after growth overnight. Infection was stopped and the bacteria were separated from the supernatant by centrifugation at 13,000 rpm for 1 min at 4°C. The cells were lysed by resuspension in 0.2 mg lysozyme and incubated at room temp for 10 minutes. Viral RNA, from both cells and supernatant, was extracted as described above (MS2 RNA isolation). Illumina NGS library was prepared and analyzed as described above.

### Haplotype inference for intracellular replication assay

To generate the replication assay plot, we had to infer haplotype data despite the fact that the populations were sequenced using short-read Illumina sequencing. However, we were able to rely on LoopSeq synthetic long reads from previous experiments (Figures 2B and 4B). In the MOI 10 experiment, each of the green helper mutations was never found one with each other, and almost always appeared in combination with the A1744G-pink mutation, allowing us to assume the same behavior during the replication assay. In the MOI 1 experiment, we made the following assumptions: (A) Mutation C1549U (light purple) in line 37B only appeared as part of a quadruplet with the other purple mutations and A1664G-orange, so we used its frequency for the entire group and reduced the frequencies of individual mutations accordingly. (B) Mutation C2859U (purple) appeared either as a pair with A1664G-orange or as a triplet with C1718U (dark purple) and A1664G-orange at similar frequencies; we divided C2859U’s remaining frequency equally between the two. We then adjusted C1718U and A1664G-orange frequencies by subtracting the triplet and pair frequencies, respectively. (C) The remaining frequency of C1718U was assumed to be in combination with A1664G-orange. (D) At MOI 1, mutation A535G-turquoise was present at no more than 5%, with the remainder assumed to occur on the same haplotype as A1664G-orange. At MOI 0.01, from 0 to 60 minutes, the same haplotype structure as in MOI 1 was assumed. At 90 minutes, the pair of A1664G-orange and A535G-turquoise remained at the same frequency as at 60 minutes. At the ON timepoints, A535G-turquoise was considered to be 50% on its own and 50% in combination with A1664G-orange. This approach allowed us to piece together short Illumina reads into coherent haplotypes, ensuring that the replication assay accurately reflected the underlying genetic structure.

### Intracellular replication model

We present a mathematical model that describes the intracellular replication cycle, encompassing the stages of entry, replication, and lysis. The lysis stage is assumed to capture both the packaging and exit processes. Additionally, we introduce a re-infection stage to account for re-infection at low MOI. Each parameter in the model (Table 3) is assumed to affect the frequency of specific genotypes at various stages of intracellular replication. Thus, for example, more rapid replication of a mutant would lead to increased frequency of this mutant at times points where replication occurs (15 through 60 mins; Methods, Fig. 1). We next used an approximate Bayesian computation (ABC) (Beaumont et al., 2002) approach to infer parameters for specific genotypes.

**Table 3.** Parameters *ω_s,m_* of the intracellular replication model.

|  | *orange + purple combo* | *orange + turquoise* | *turquoise* | *orange* | *wt* |
| --- | --- | --- | --- | --- | --- |
| *entry* | $\omega_{e,oc}$ | $\omega_{e,tu}$ | $\omega_{e,tu}$ | 1 | 1 |
| *replication* | $\omega_{r,oc}$ | 1 | 1 | 1 | 1 |
| *lysis* | $\omega_{l,o} + \delta$ | $\omega_{l,o}$ | 1 | $\omega_{l,o}$ | 1 |
| *re – infection* | $\omega_{rei,o}$ | $\omega_{rei,o}$ | $\omega_{rei,tu}$ | $\omega_{rei,o}$ | 1 |

Let *ω_stage,mutation_* ∈ ℝ^+^ be the parameters of our model, where the stage is denoted as *s* ∈ {*entry* (*e*), *replication* (*r*), *lysis* (*l*), *re* − *infection* (*rei*)} and the mutations (haplotypes) as *m* ∈ {*A*1664*G*, *A*1664*G and purble helper mutations*, *A*535*G*, *A*1664*G and A*535*G*, *wt*}.

We define: *o* (*orange*) ≔ *A*1664*G*, *oc* (*orange* + *combo*) ≔

*A*1664*G and purble combo helper mutations*, *tu* (*turquoise*) ≔ *A*535*G*,

*otu* (*orange* + *turquoise*) ≔, *A*1664*G and A*535*G*.

Hence, *m* ∈ {*o*, *oc*, *tu*, *otu*, *wt*}.

Each parameter *ω_s,m_* represents the “fitness” of the haplotypes *m* at the stage *s* of the replication cycle. We also introduce *δ* ∈ ℝ as the parameter representing the added delay in lysis due to the presence of the combination of the purple helper mutations over the orange mutation.

To avoid over-parameterization, we made the following assumptions:

1. Parameters for haplotypes that do not affect certain stages of the replication cycle, were fixed at 1. Thus, for example, a non-synonymous mutation in the lysis protein was assumed to have no impact on entry. Accordingly, entry parameters were assumed only for non-synonymous mutations at A or coat proteins. Replication parameters were assumed only for non-synonymous mutations at the replicase protein, mutations that affected the MJ structure or the CT structure, or mutations affecting the coat protein that interacts with the TR loop. Lysis parameters were assumed for non-synonymous mutations at the lysis protein or mutations affecting the LH structure.
2. For haplotypes consisting of multiple mutations, the parameters were determined by the mutation most relevant to the specific stage of the replication cycle. For example, the orange mutation (A1664G) affects the lysis, therefore *ω_l,otu_* = *ω_l,o_*.

### Simulation of haplotype frequencies

We simulated the frequencies of the observed haplotypes at each time point in our second intracellular replication assay (based on the original passaging experiment at MOI=1) which was conducted at MOI of 0.01 (Fig. 5A). The objective was to infer the effects of individual mutations or combinations of mutations on the different stages of the replication cycle. The set of time points is defined as *T* = {*t*_2_, *t*_34_, *t*_52_, *t*_64_, *t*_72_, *t*_84_, *t*_92_}, where frequencies are simulated at each time point. Notably, *t*_84_ is simulated solely as a correction for uniform time intervals of 15 minutes, with no data fitting at that time point, as no empirical data exists for it.

Based on Fig. S6, we define indicator vectors *I_t_* ∈ {0,1}^4^ which indicate the active stages - entry, replication, lysis and re-infection respectively, at each time point, as follows:

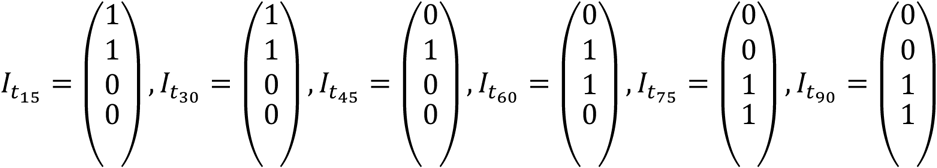

Uncorrected haplotype frequencies at a given time *t* are given by:

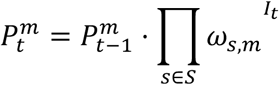

To compute the corrected frequencies, we normalize the haplotype frequencies by calculating the relative abundance:

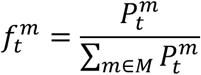

Initial empirical frequencies at time point zero *t*_2_ were used to initialize the simulations.

### Inference of model parameters

We employed a wide prior for the “fitness” parameters thus minimizing prior assumptions and allowing for a broad range of parameter sampling, with *ω_s,m_* ∼ *U*(0,4) *for all s* ∈ *Stages and m* ∈ *Mutations*. For the additional lysis delay parameter, representing the possible effect of the purple helper mutations (denoted as combo), we used a prior *δ*∼*U*(−1,1), aiming to assess whether these helper mutations contribute additional delay beyond the orange mutation (A1664G) alone.

Given the complexity and large number of parameters, traditional maximum likelihood estimation was impractical. Therefore, we utilized approximate Bayesian computation with sequential Monte Carlo (ABC-SMC) (Lintusaari et al., 2017), as implemented in the Python package pyABC (Klinger et al., 2018). As a summary statistic we used the sum of squared differences between the simulated and empirical data across all time points, calculated using the following squared ℓ_2_ distance function:

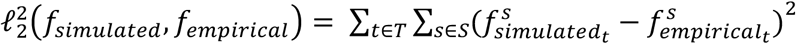

The ABC-SMC algorithm was run with threshold *ε* that decreased to 0.078 over 15 iterations. Each iteration used 10,000 particles, and the particles of the final iteration were used to generate posterior distributions for all parameters.

## Data and code availability

All sequencing data presented in this paper are available in the sequencing read archive (SRA). Previously published data (MOI=1 passages 1 though 23) are available under under BioProjects PRJNA575138, PRJNA547685. All other sequencing data are available under BioProject PRJNA1161593. Processed data including inferred mutation frequencies and haplotypes are available in the Zenodo database under accession code 10.5281/zenodo.13764969. All code used for sequence analysis and for modelling and parameter inference is available at https://github.com/Stern-Lab/cheaters_fine_balance.

## Acknowledgements

We thank Avigdor Eldar and Danielle Miller for stimulating conversations and insightful reading. This study was supported by an ERC starting grant 852223 (RNAVirFitness) to AS and by a European Research Council grant AdG 787514 to UG. This study was also supported by a fellowship to Y.M and A.BZ from the Edmond J. Safra Center for Bioinformatics at Tel Aviv University.

## Notes

### Competing Interest Statement

The authors have declared no competing interest.

