## Supplementary information for "Navigating a fine balance: point-mutant cheater viruses disrupt the viral replication cycle"

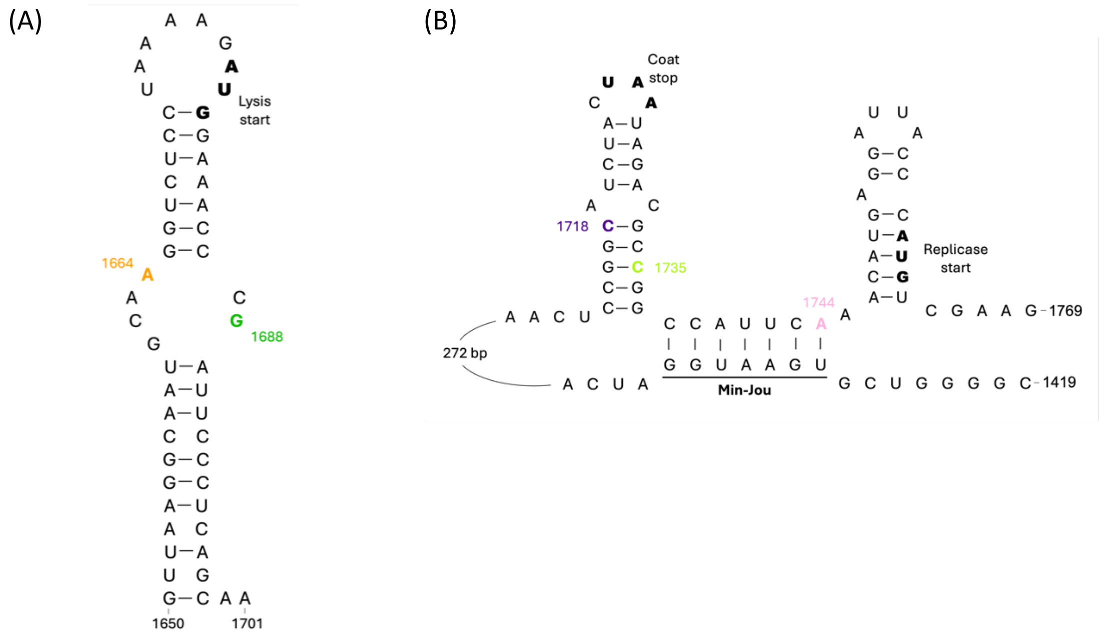


**Figure S1. RNA secondary structures in MS2 together with locations of mutations found herein that impact these structures.** (A) Lysis hairpin. (B) A regulatory structure including the coat gene terminator (CT) loop (left), Min-Jou (MJ; center) and TR loop (right). Previous work has shown that mutations at the CT as well as in the MJ affect replicase translation.

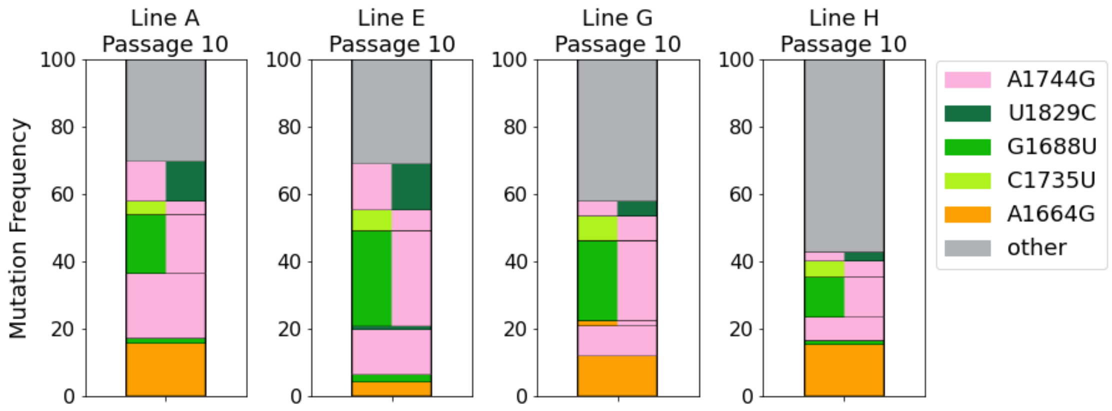


**Figure S2. Haplotype analysis of parallel evolution during MOI=10.** Figure similar to Fig. 2B with a lower frequency threshold of 1%. Green mutations are never found together and very rarely on their own.

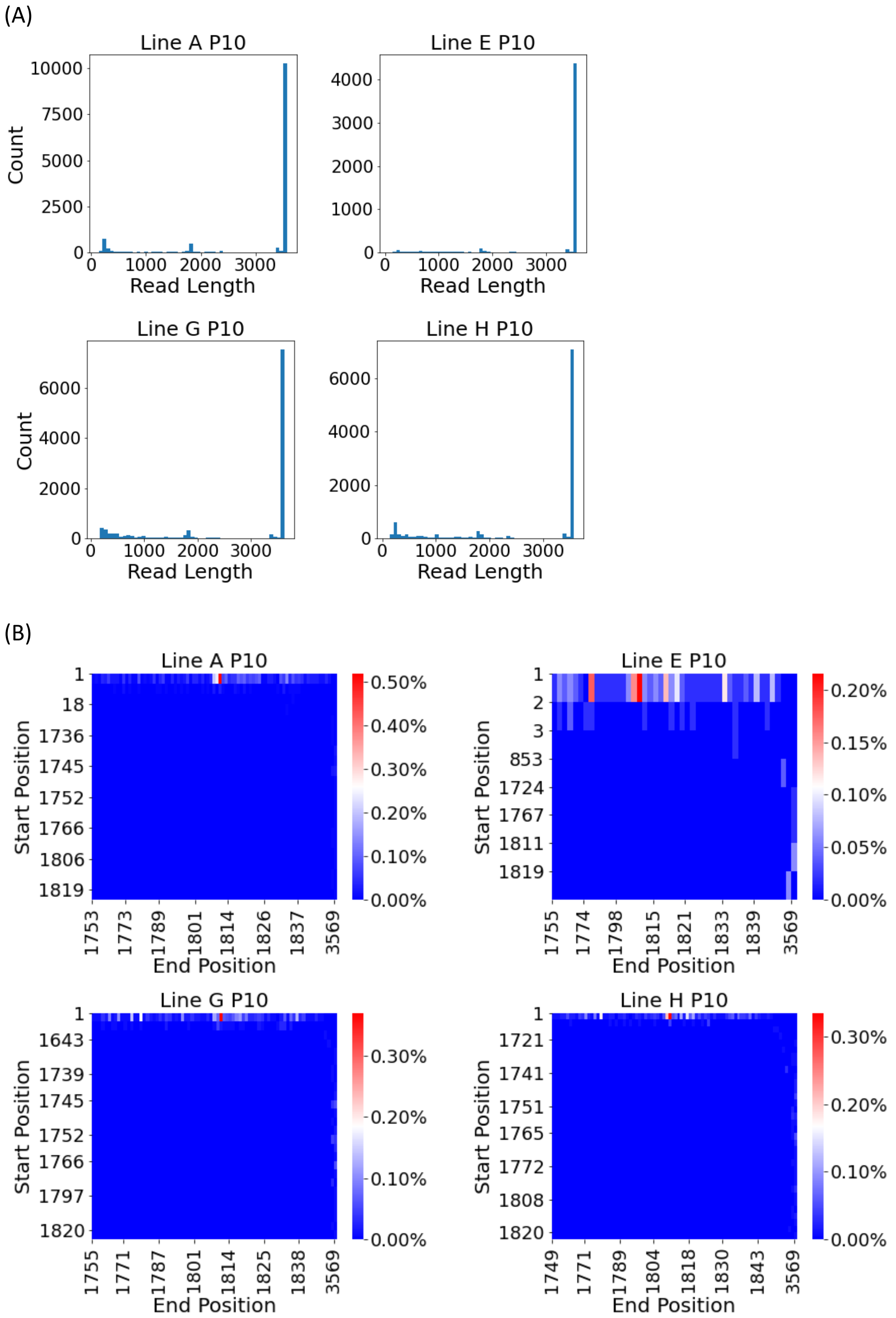


**Figure S3. Analysis of short genome fragments in synthetic long-read sequencing of parallel evolution during MOI=10.** (A) The distribution of the read lengths obtained in synthetic long-read sequencing of p10. The vast majority of reads are full length genomes, a small minority of short genomic fragments is observed, with a consistent peak over ~1800 base long fragments. (B) Heatmap illustrating the frequency of reads across each start and end position, only fragments ranging between 1750 and 1850 nucleotides in length are shown. While there is presence of short genomic fragments, their frequency at passage 10 does not exceed 0.5%.

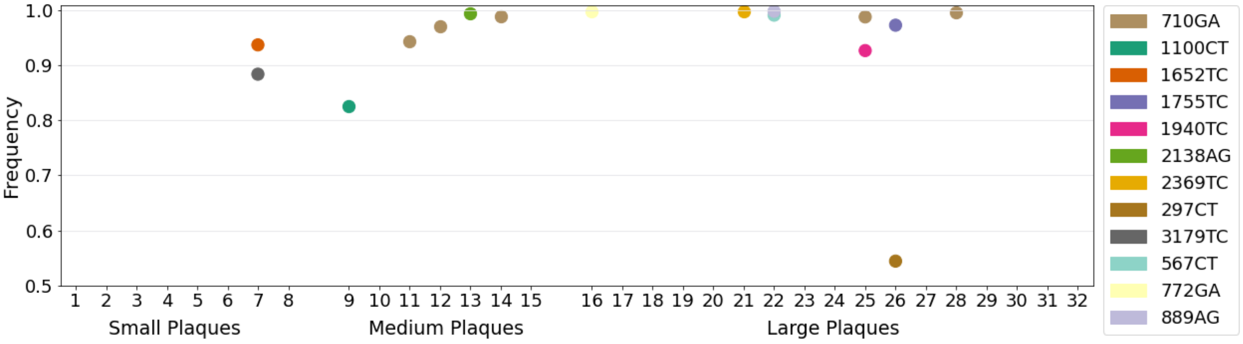


**Figure S4. Sequencing of plaques in various sizes.** Mutation frequencies found in 32 plaques of various sizes (small, medium and large) isolated from line p8-D, MOI = 10 experiment. Mutations exceeding 50% are shown. Plaques were numbered, originally 40 plaques were isolated but some were not sequenced successfully.

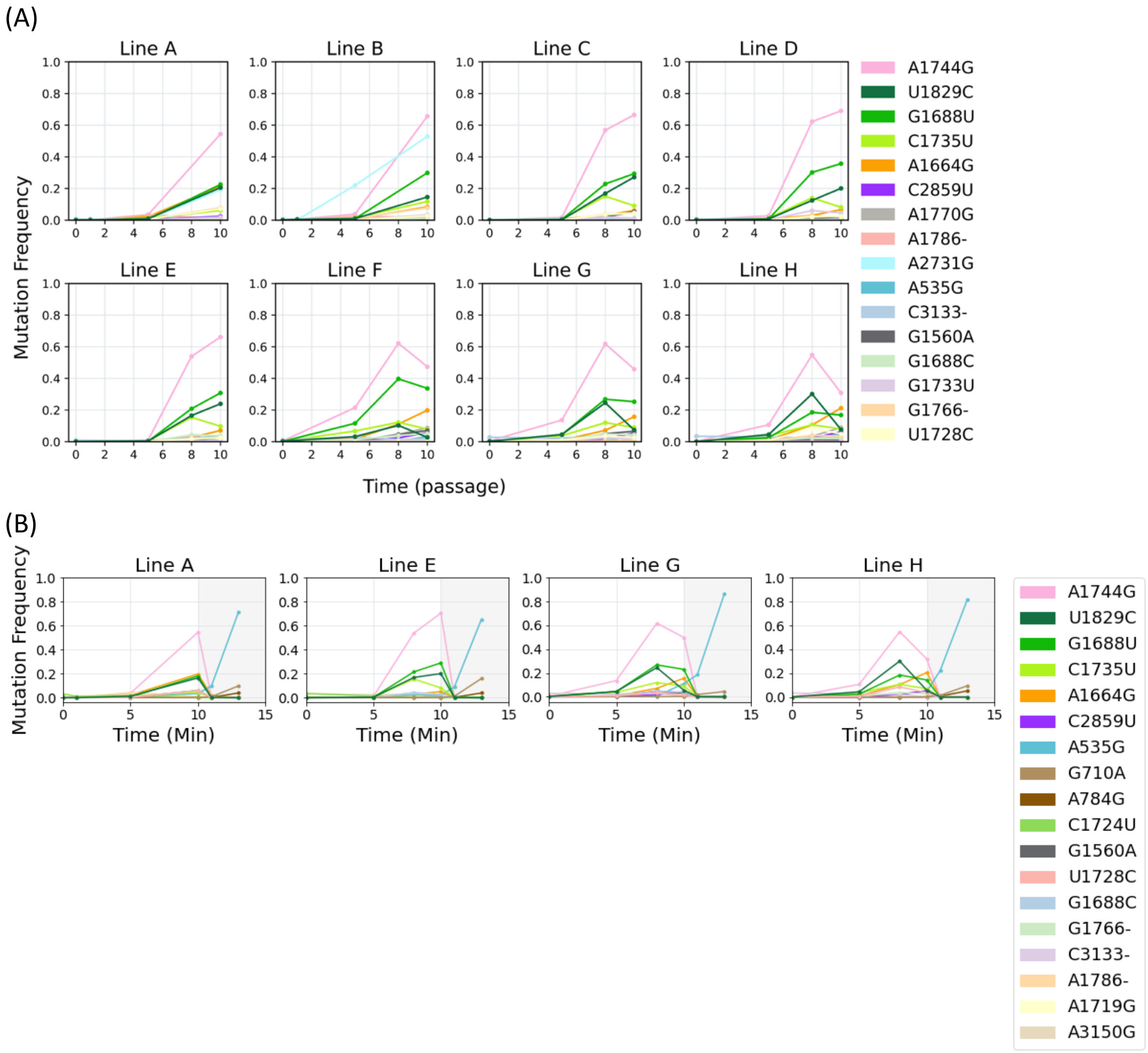


**Figure S5. Parallel evolution during MOI=10 passaging. Figure corresponds to figure 2 with a lower frequency threshold of 3%.** (A) Corresponds to Fig. 2A (B) Corresponds to Fig. 2C.

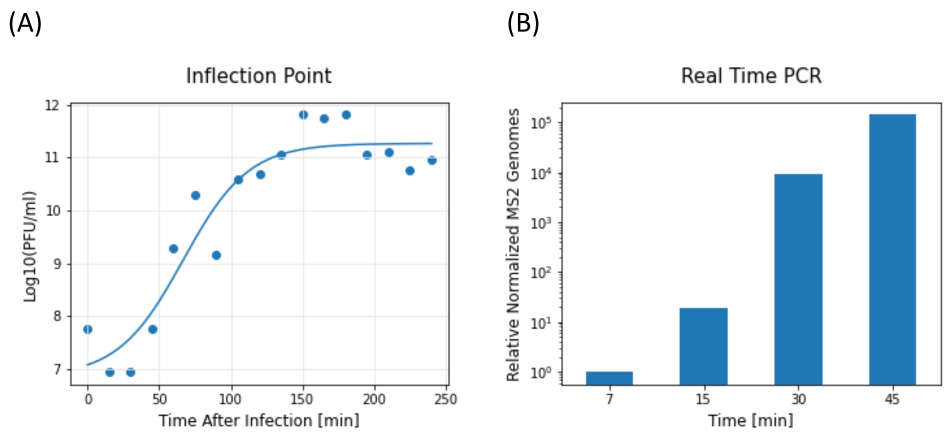


**Figure S6. MS2 virus replication cycle measurement at MOI 1.** Taken from (Meir et al., 2020). (A) Assessment of the viral replication cycle by measuring time to cell lysis. Particle-forming units (PFU) per ml were recorded at various intervals over a 250-minute period, allowing us to pinpoint the inflection point. This experiment, performed with WT viruses following the procedures outlined in the Methods, revealed that 50% of bacterial cells lysed approximately 67 minutes after infection (inflection point of the graph). (B) Intracellular viral RNA levels were measured at 7, 15, 30, and 45 minutes post-infection. Cells infected with WT viruses underwent RNA quantification via qPCR, with normalization to the 16S gene. The RNA levels at each time point were expressed relative to the 7-minute mark, post-adsorption. Replication was observed throughout all times tested, as evident by the increase in RNA levels.

```
Query ID: AGD81189.1 (Qbeta replicase protein)

Sbjct: replicase [Escherichia phage MS2]
Sequence ID: ABQ02458.1Length: 545Number of Matches: 1

Query  68    IDYLKAEIMSKYGDFSLGIDT---EAVAWEKFLAAEAECALTNARLYRPDYSEDFNFSLG  124
             I YL+ E+++K+     G D     A+A  K   A   C   N    R  +  D + S
Sbjct  45    IAYLRDELLTKHPSLGNGNDEATRRALAIAKLREANERCGQIN----REGFLHDKSLSWD  100

Query  125   ESCIHMARRKIAKLIGDVPS--VEGMLRHCRFSGGATTTNNRSYGHPSFKFALPQACTPR  182
                +  +   I  LIG++ S     +   C FS GA+  +      P  KFA     TPR
Sbjct  101   PDVLQTS---IRSLIGNLLSGYRSSLFGQCTFSNGASMGHKLQDAAPYKKFAEQATVTPR  157

Query  183   ALKYVLALRAS----THFDIRISDISPF-----NKAVTVPKNSKTDRCIAIEPGWNMFFQ  233
             AL+  L +R          +R ++   F     N   TVPKN+K DR    EP  NM+ Q
Sbjct  158   ALRAALLVRDQCAPWIRHAVRYNESYKFRLVVGNGVFTVPKNNKIDRAACKEPDMNMYLQ  217

Query  234   LGIGGILRDRLRCWGIDLNDQTINQRRAHEGSVTNNLATVDLSAASDCISLALCELLLPP  293
              G+G  +R RLR  GIDLNDQTINQR A +GSV  +LAT+DLS+ASD IS  L    LPP
Sbjct  218   KGVGAFIRRRLRSVGIDLNDQTINQRLAQQGSVDGSLATIDLSSASDSISDRLVWSFLPP  277

Query  294   GWFEVLMDLRSPKGRLPDGSVVTYEKISSMGNGYTFELESLIFASLARSVCEILDLDSSE  353
               +  L  +RS  G + DG  + +E  S+MGNG+TFELES+IF ++ ++  +I   ++
Sbjct  278   ELYSYLDRIRSHYG-IIDGETIRWELFSTMGNGFTFELESMIFWAIVKAT-QIHFGNAGT  335

Query  354   VTVYGDDIILPSCAVPALREVFKYVGFTTNTKKTFSEGPFRESCGKHYYSGVDVTPFYIR  413
             + +YGDDII PS   P + E   Y GF  N +KTF  G FRESCG H+Y GVDV PFYIR
Sbjct  336   IGIYGDDIICPSEIAPRVLEALAYYGFKPNLRKTFVSGLFRESCGAHFYRGVDVKPFYIR  395

Query  414   HRIVSPADLILVLNNLYRWATIDGVWDPRAHSVYLKYRKLLPKQL  458
               + +   L+L++N L  W  + G+ DPR + V+++   L+P
Sbjct  396   KPVDNLFSLMLIMNRLRGWGVVGGMSDPRLYKVWVRLSSLVPSMF  440
```

**Fig. S7. Local alignment of Qbeta replicase protein (Query) and MS2 replicase protein (Sbjct) obtained via BLAST** (Altschul et al., 1990; McGinnis & Madden, 2004). Shown in red is motif E, one of the five motifs shared among most RNA-dependent-RNA polymerases (Gong, 2021) which in principle interacts with product RNA. Site 376 in MS2 rep is site 394 in qbeta replicase, one amino acid upstream of motif E and marked in bold.

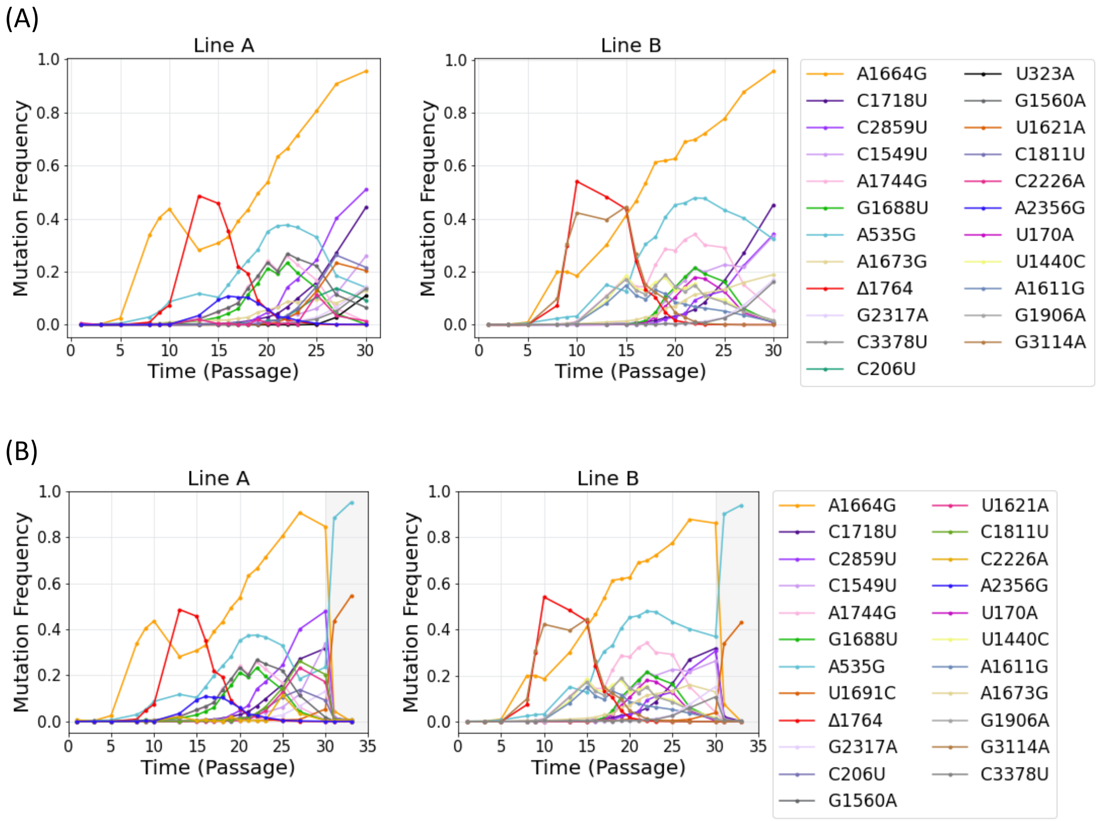


**Figure S8. Serial passaging experiments at the MOI of 1.** Similar to Fig. 4A,C, with lower frequency threshold of 3%.

(A)

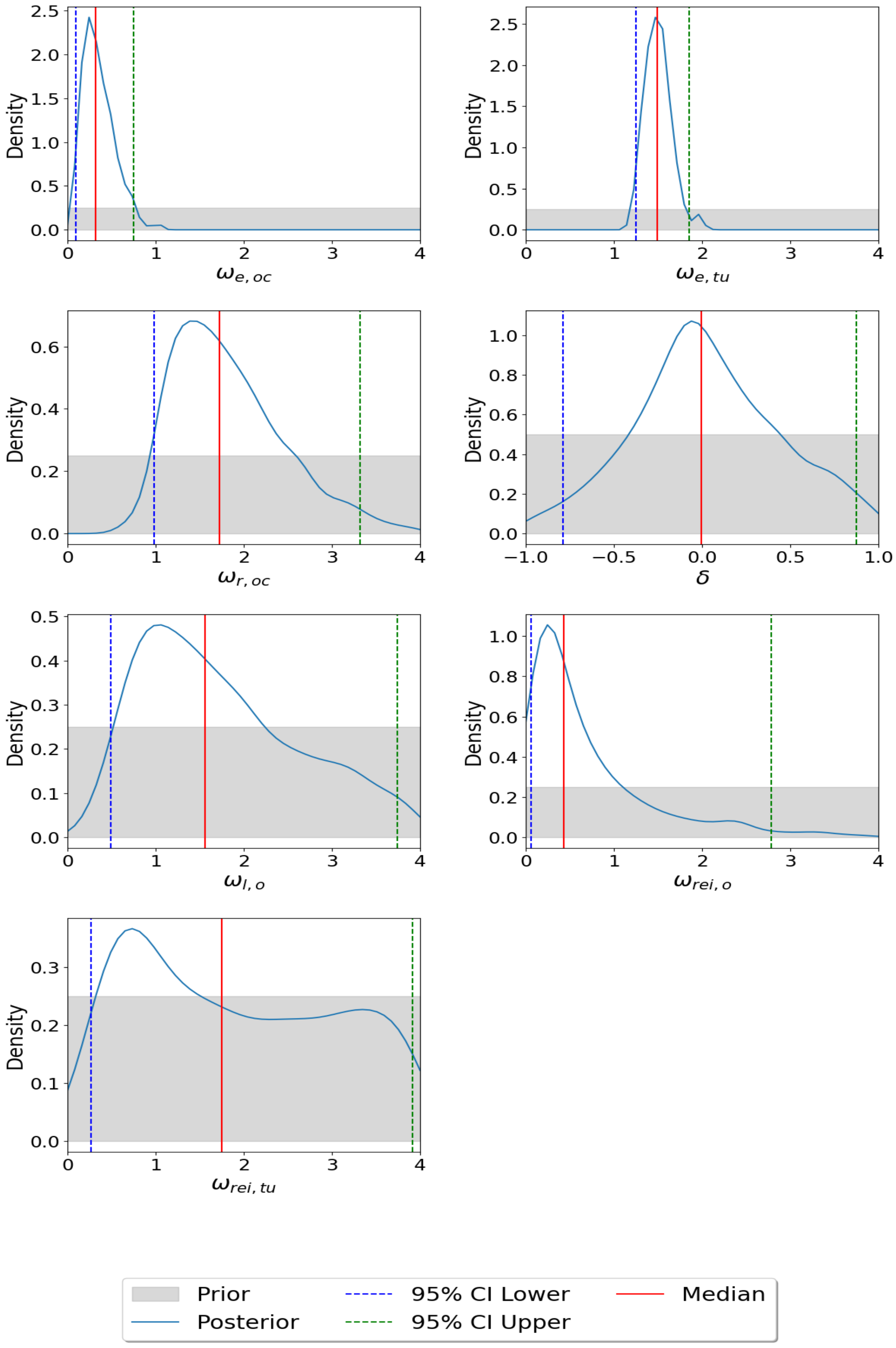

Density
$\omega_{e,oc}$
$\omega_{e,tu}$
$\omega_{r,oc}$
$\delta$
$\omega_{l,o}$
$\omega_{rei,o}$
$\omega_{rei,tu}$
Prior
Posterior
95% CI Lower
95% CI Upper
Median

(B)

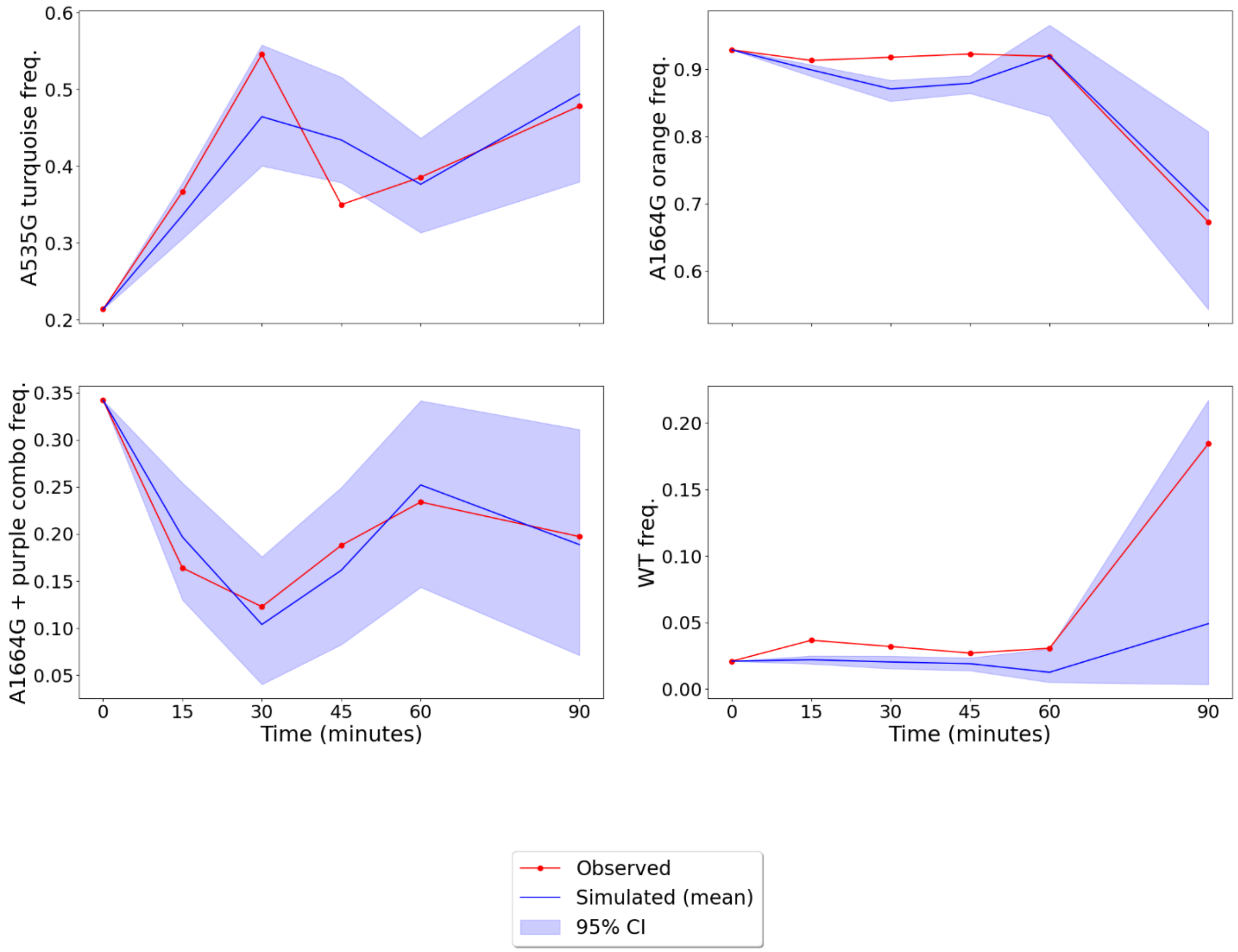


**Figure S9. Inferences of parameters from intracellular replication model.** (A) Marginal posterior distributions for all parameters of the model. The red line shows the median value, and the dashed lines are the bounds of the 95% highest density interval (HDI), marked as upper or lower confidence intervals (CIs). The grey background illustrates the uniform prior distribution. (B) Posterior predictive checks showing the empirical (observed) data in red versus simulated data using parameters from the 95% HDI of the posterior (light blue band), average value shown in dark blue.

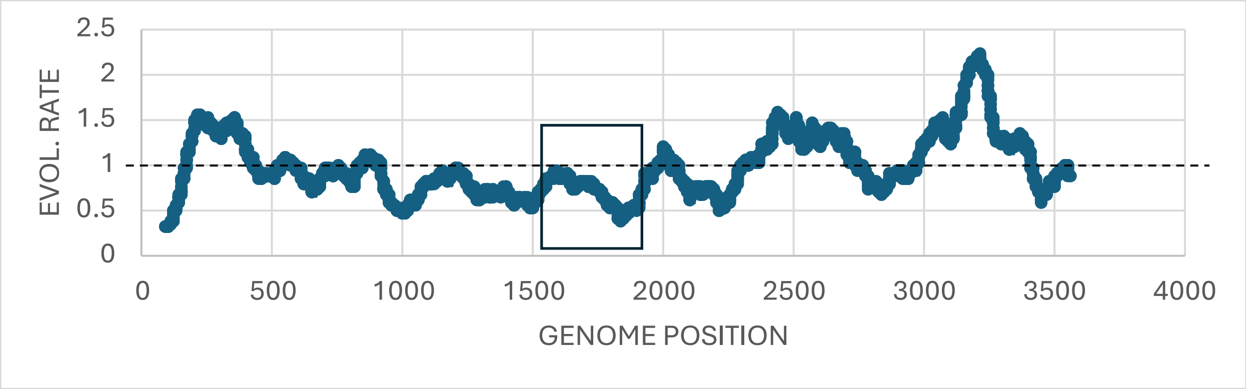


**Figure S10. Site specific evolutionary rate of MS2 along the genome.** We began by performing BLAST (Altschul et al., 1990; McGinnis & Madden, 2004) of the MS2 genome against the nr NCBI database. This yielded for the most part synthetic constructs that were removed from the analysis. We continued by searching for relatives of MS2 in a recently published paper that analyzes metagenomic data to expand the RNA virome (Neri et al., 2022). We focused only on relatives of MS2 in the Fiersviridae tree with a distance along the tree lower than 0.5, to avoid very distant relationship. We further added three phages discovered in a more recent such scan (Quinones-Olvera et al., 2024), leading to a total of 17 homologous sequences. Sequences were aligned using MAFFT (Katoh & Standley, 2013) and given as input to PhyML (Guindon et al., 2005) for maximum-likelihood phylogenetic inferences; both programs were run with default parameters. The resulting phylogeny and alignment were used as input for rate4site (Mayrose et al., 2004; Pupko et al., 2002) for inference of site-specific rates. The figure shows values of average rates over sliding windows of size 100. The average rate of 1 is shown by a dashed horizontal line. The region where most cheater and helper mutations were found (main text) is boxed.

**Table S1. MS2 primers for NGS library preparation**

| Primer | Sequence |
|---|---|
| 3-F1 (F) | GGGTGGGACCCCTTTCGG |
| 3-R1 (R) | GCTAACGCATCTAAGGTATGG |
| 3-F2 (F) | GGCCCAAATCTCAGCCATGC |
| 3-R2 (R) | CGTGTCTGATCCACGGC |
| 3-F3 (F) | GGCACAAGTTGCAGGATGCA |
| 3-R3 (R) | TGGGTGGTAACTAGCCAAGCAG |
| 7-F1 (F) | GGGTGGGACCCCTTTCGG |
| 7-R1 (R) | CTTCACGAGCGCAATGGTTTG |
| 7-F2 (F) | GCAAAAGGTCACCCAGGGTAAT |
| 7-R2 (R) | ATGATGGACTCACCCGTTATTACG |
| 7-F3 (F) | CTGTAGGTAACATGCTCGAGGG |
| 7-R3 (R) | AGCGAAAATTGGAATGGTTAGTTCC |
| 7-F4 (F) | AAAGTCGAGGTGCCTAAAGTGG |
| 7-R4 (R) | CCAGAGAGGAGGTTGCCAATAAG |
| 7-F5 (F) | TGATCGGTGCGGTCAGATAA |
| 7-R5 (R) | TGAATATAGCTCAGGTGGGAGAAAAC |
| 7-F6 (F) | GATGGTTCGCTTGCGACGATA |
| 7-R6 (R) | CACCGAAGAACATCGAAGGCA |
| 7-F7 (F) | TGTTGACAATCTCTTCGCCCTG |
| 7-R7 (R) | TGGGTGGTAACTAGCCAAGCAG |
